# Phenotypic and Developmental Dissection of an Instance of the Island Rule

**DOI:** 10.1101/2025.01.22.634334

**Authors:** Mark J. Nolte, Bret A. Payseur

## Abstract

Organismal body weight correlates with morphology, life history, physiology, and behavior, making it perhaps the most telling single indicator of an organism’s evolutionary and ecological profile. Island populations provide an exceptional opportunity to study body weight evolution. In accord with the “island rule,” insular small-bodied vertebrates often evolve larger sizes, whereas insular large-bodied vertebrates evolve smaller sizes. To understand how island populations evolve extreme sizes, we adopted a developmental perspective and compared a suite of traits with established connections to body size in the world’s largest wild house mice from Gough Island and mice from a smaller-bodied mainland strain. We pinpoint 24-hour periods during the third and fifth week of age in which Gough mice gain exceptionally more weight than mainland mice. We show that Gough mice accumulate more visceral fat beginning early in postnatal development. During a burst of weight gain, Gough mice shift toward carbohydrates and away from fat as fuel, despite being more active than and consuming equivalent amounts of food as mainland mice. Our findings showcase the value of developmental phenotypic characterization for discovering how body weight evolves in the context of broader patterns of trait evolution.

## Introduction

Organismal body size correlates with morphology, life history, physiology, and behavior (Peters, 1983; Calder, 1984; Schmidt-Nielsen, 1984; Bonner, 2024). As a result, body size is perhaps the most telling single indicator of an organism’s evolutionary and ecological profile. Expansive interest in body size has stimulated a search for general patterns that characterize its evolution. For example, body size increases with latitude (Bergmann, 1847), shows a right skew in its distribution (Hutchinson & Macarthur, 1959), and displays a bias toward increasing over geological time (Simpson, 1944) (Cope’s rule) across species of mammals and other vertebrates.

One of the more striking observations pertaining to the evolution of body size is captured by the “island rule.” On islands, small-bodied vertebrates tend to become bigger and large-bodied vertebrates tend to shrink (Foster, 1964; Van Valen, 1973; Lomolino, 1985, 2005; Benítez-López et al., 2021). Comparisons between island and mainland species point to potential ecological causes of this intriguing biogeographic pattern, including substantive changes in predation and competition for resources (Heaney, 1978; Adler & Levins, 1994; Raia & Meiri, 2006; Lomolino et al., 2012; Durst & Roth, 2015; Cuthbert et al., 2016; Itescu et al., 2020; Benítez-López et al., 2021; Ponti et al., 2023).

Understanding how island populations achieve extreme body sizes ultimately requires in-depth characterization of organismal growth. Evolutionary developmental biology and evolutionary quantitative genetics provide a strong conceptual foundation for this approach (Atchley et al., 2000). Broadly speaking, evolutionary changes in body size are expected to be accomplished by age-specific shifts in genetic regulation, cellular processes, and endocrinological factors (Atchley et al., 2000; Cheverud, 2005; Kenney-Hunt et al., 2006; Wilches et al., 2021). Documenting and quantifying shifts in these “intermediate phenotypes” or “endophenotypes” between populations accomplishes two primary functions. First, by pinpointing endophenotypic shifts, specific organismal characteristics shaped by natural selection and other evolutionary forces are revealed, yielding clues into how these forces operate. Second, the discovery of “endophenotypes” holds the promise of connecting a downstream composite phenotype of interest (e.g. body weight) with its genetic causes (Landry & Rifkin, 2012; Rittschof & Robinson, 2016). Despite the benefits of this framework, the developmental, physiological, and morphological correlates of extreme body size are rarely examined in island populations.

Rodents are promising subjects for discovering the organismal processes that underlie body size evolution on islands. The repeated and convergent evolution of large size in island rodents provides some of the strongest evidence for the island rule (Foster, 1964; Redfield, 1976; Lomolino, 1985, 2005; Adler & Levins, 1994; Case et al., 2002; Meiri et al., 2008; Durst & Roth, 2015; Nolfo-Clements et al., 2017; Baier & Hoekstra, 2019; Benítez-López et al., 2021). Classic research focused on the ecology and evolution of house mice uncovered multiple examples of island gigantism within a single species (Berry, 1964; Berry & Peters, 1975; Berry et al., 1978a,b, 1979; Rowe-Rowe & Crafford, 1992). In addition, house mice can be raised in the lab, making it possible to characterize evolved genetic differences between island and mainland populations.

The world’s largest wild house mice inhabit remote Gough Island (Rowe-Rowe & Crafford, 1992; Gray et al., 2015), located in the South Atlantic Ocean. Mice colonized Gough Island from Western Europe (Wace, 1961) a few hundred generations ago (Rowe-Rowe & Crafford, 1992; Gray et al., 2014), suggesting the evolution of extreme size was rapid. Genetic mapping identified quantitative trait loci (QTL) that contribute to a striking body size difference between Gough Island (Gough) mice and mice from a mainland strain (Gray et al., 2015), as well as QTL for enlarged skeletal elements (Parmenter et al., 2016) and increased skeletal load resistance (Payseur et al., 2023). Comparisons between Gough mice and mainland mice uncovered differential expression at thousands of genes in metabolic organs that could be connected to weight accumulation (Nolte et al., 2020). The application of genetic and molecular tools available for biomedical and evolutionary studies of house mice have established Gough mice as a powerful model for the genetic dissection of the island rule and mammalian growth.

To gain deeper insights into the genesis of a single instance of the island rule, we characterize and compare a wide range of endophenotypes with established connections to body size in Gough mice and mainland mice. We document characteristics of island-mainland divergence in body weight and growth from birth to six weeks of age, with temporal resolutions ranging from 24-hour periods to weekly intervals. We report divergent endophenotypes that span numerous levels of biological organization, including organ weights, circulating hormones, cellular dynamics, and metabolic fuel processing. Our comparative investigation of growth at an unusually high resolution is a promising first step toward elucidating causes of the island rule.

## Materials and Methods

### Mice and mouse husbandry

Two inbred strains of the Western European house mouse (*Mus musculus domesticus*) were used in this study. Gough Island mice, the larger of the two strains, were sourced from a laboratory colony derived from wild live-caught mice from Gough Island that were shipped to the University of Wisconsin School of Veterinary Medicine Charmany Instructional Facility in 2009 (Gray et al., 2015). WSB/Eij is a wild-derived, inbred strain with a body size representative of mainland wild house mice; those used in this study derived from mice purchased from The Jackson Laboratory (Bar Habor, ME) in 2009. All mice were kept in a temperature-controlled room (20-22.2 °C/68–72 °F) set on a 12-hour light/dark cycle (6 am/6 pm). All pups were weaned at 3 weeks and housed with one to three other individuals of the same sex. Weaned mice were housed in micro-isolator cages with corn cob substrate (0.32 cm; The Andersons Lab Bedding) and *ad libitum* access to water and food (Envigo 2020X Teklad Global Diet). Mice paired for mating and their pups were provided with a higher fat chow (Envigo 2019 Teklad Global Diet) and an igloo (Bio Serv). All protocols were approved by the University of Wisconsin-Madison Institutional Animal Care and Use Committee in compliance with the US National Research Council’s Guide for the Care and Use of Laboratory Animals, the US Public Health Service’s Policy on Humane Care and Use of Laboratory Animals, and Guide for the Care and Use of Laboratory Animals.

### Daily body weights

Body weights were taken daily between 8 am and 11 am from the day of birth (day zero) to day 42. Mice were momentarily placed in a plastic container and weighed on a Torbal AGC100 Lab Scale (100g x 0.001g). The Torbal AGC100 Lab Scale was set to “Animal weigh” mode.

### Organ weights

Prior to organ (and plasma) collection, mice were fasted for an average of 5 hours and 33 minutes. Mice were euthanized using non-vaporized isoflurane calibrated to mouse weight and cervical dislocation. Organs were weighed using a Torbal AGN100 Internal Calibration Analytical Balance, 100 g x 0.0001 g, after being collected in the following order: pancreas, mesenteric fat, liver, gonadal fat, retroperitoneal fat, inguinal subcutaneous fat, scapular brown fat, quadriceps muscle, triceps surae muscle(s), brain and intestine (length in cm). To reduce desiccation, collection was completed within 24.5 minutes (SD = 1.4 minutes), on average.

For each organ, between 13 and 15 samples were collected. For each organ and week, differences in organ weight between Gough and mainland mice were determined using t-tests. P-values were assessed against a range of critical values that correct for familywise error rates (Bonferroni, Holm, and Benjamini-Hochberg).

Determinants of organ weight were evaluated using factorial ANOVA with sex, strain, and week as main effects, and a strain-by-week interaction. To ensure homoscedasticity and normality of the residuals, weights for liver, mesenteric fat, pancreas, and triceps surae were log-transformed; weights for gonadal fat and retroperitoneal fat were square-root transformed. For each organ and weekly transition, differences in growth rate between strains were determined using nonorthogonal contrasts that tested whether the change in mean organ weight from an earlier week to a later week (the slope) was different between the strains. P-values were assessed against a range of critical values that correct for familywise error rates (Bonferroni, Holm, and Benjamini-Hochberg).

### Blood plasma molecules

Blood was collected from fasted mice. Fasting began in the morning between 8 am and 10:30 am. Prior to collection, mice were anesthetized using non-vaporized isoflurane. Using heparin-coated capillary tubes, blood was collected via retro-orbital blood draw into 0.02% EDTA-coated tubes placed on ice for 1-5 hours. Tubes were centrifuged at 1300 RPM for 10 minutes and retrieved plasma was stored at −80° C.

Ten samples per condition (sex/strain/week) for each of eight plasma analytes with established connections to body size were quantified at the University of Wisconsin Primate Center: adiponectin: a singleplex Bio-Rad Bio-Plex Pro Mouse Luminex immunoassay kit (cat. no. 171F7002M); insulin, ghrelin, glucagon, and leptin: a multiplex Bio-Rad Bio-Plex Mouse Diabetes Panel Luminex immunoassay kit (cat. no. 171F2001M); IGF-1: an R&D Systems Mouse/Rat Quantikine IGF-1 ELISA kit (cat. no. MG100); free fatty acids: a Millipore-Sigma Free Fatty Acid Quantification Kit (cat. no. MAK044; the calorimetric protocol was followed using 5 ul plasma samples in 96-well plates); glucose: a YSI 2600 biochemistry analyzer using 13 ul plasma samples. Quantification was done in duplicate. Outliers were removed in two steps. First, for each analyte and condition the absolute value of the difference between duplicate values was box plotted. Second, the midpoint value for accepted samples was box plotted. For both steps, removed outliers fell outside the interquartile range multiplied by 1.5 and added to quartiles 1 and 3.

Determinants of plasma analyte concentration were evaluated using ANOVA with strain and week as the main explanatory effects, and a strain-by-week interaction. To ensure homoscedasticity and normality of the residuals, concentrations for adiponectin and leptin were log-transformed. For each plasma analyte, differences between strains were determined using t-tests. Resulting p-values were assessed against a range of critical values that correct for familywise error rates (Bonferroni, Holm, and Benjamini-Hochberg).

### Cell proliferation

The liver, gonadal fat pad, and femur were collected at three periods: day 15 (both sexes); day 30 (females); day 32 (males). The left lobe of the liver was collected. Left and right female gonadal fat was taken from the border between the parametrial and perivesical portions of the abdominopelvic fat depot (Vitali et al. 2012). In males, the distal half of both testicular fat depots was collected.

All tissues were fixed in 10% neutral buffered formalin for 24 hours at room temperature, washed three times in 1X PBS for 10 minutes, moved into 70% ethanol, and stored at 4° C prior to paraffin embedding. Day 15 and day 30 or 32 samples were stored at 4 C for at least three and seven days, respectively. Prior to the three PBS washes, approximately 1/3 of the right and left margins of the liver were cut away and a cut surface was embedded face-down in paraffin. For the femurs a decalcification step was added after the PBS washes. Day 15 and day 30 or 32 femurs were decalcified at room temperature for 3 and 7 days, respectively, in 0.5 M EDTA (pH 7.2), then washed three times in 1X PBS for 10 minutes.

Cell proliferation was detected using two antibodies against mitotic cell division markers, Ki-67 and phospho-histone H3 (phh3): Abcam: Ki-67 - Anti-Ki67 (ab15580); Phh3 - Recombinant Anti-Histone H3 (phospho S28) (ab32388).

Multiplex immunohistochemistry was performed on the automated Ventana Discovery Ultra BioMarker Platform after manual deparaffinization. Deparaffinization and rehydration were carried out by emerging slides in xylene twice for 5 minutes each, followed by rehydration in ethanol (100%, 95%, 70%) and distilled water for 5 minutes each. After rehydration, slides were fixed for 7 minutes in 10% formalin and then air dried before returning to the Ventana instrument for staining. Antigen retrieval and heat-induced epitope retrieval were achieved with the EDTA-based cell conditioner 1 buffer (pH 8.4; Ventana #950-224) for 48 minutes at 75 °C. The primary antibody, Phh3, was diluted 1:250 in Ventana Antibody Diluent with Casein (Roche Diagnostic Ref# 760-219) and manually added and incubated for 40 minutes at 37 ℃. Slides were rinsed with reaction buffer (Ventana #950-300), incubated with Discovery OmniMap anti-rabbit horseradish peroxidase (Ventana #760-4311) for 16 minutes at 37 ℃, and rinsed with reaction buffer (Ventana #950-300). Discovery ChromoMap DAB detection kit (Ventana #760-159) was used for visualization. Discovery Inhibitor (Roche diagnostic #760-4840) was used to denature the first antibody (without affecting signal). The secondary antibody, Ki-67, was diluted 1:100 in DaVinci Green Antibody Diluent (BioCare Medical #PD900H) for 32 minutes at 37 ℃. Slides were rinsed with reaction buffer, incubated with Discovery OmniMap anti-rabbit horseradish peroxidase (Ventana #760-4311) for 16 minutes at 37 ℃, and rinsed with reaction buffer. Discovery purple detection kit (Ventana # 760-229) was used for visualization. Upon removal, slides were counterstained with Harris hematoxylin (1:5) for 45 seconds, rinsed with dH_2_O, dehydrated by oven drying and dipping in xylene.

Stained tissue was imaged on an Olympus BX43 microscope with Perkin Elmer Nuance software (Brightfield). Images were taken at 20X (liver) and 40X (gonadal fat). Between 8 and 10 images were taken per sample.

Ki-67- and Phh3-postive cells were detected and quantified using Perkin-Elmer inForm/Nuance software after implementation of a signal-detection algorithm trained on sample images. Prior to algorithm training, the following inForm/Nuance parameters were selected: Segment Tissue – skipped; Find Features – cell segmentation; Phenotyping – skipped; Score – score. The following Cell Segmentation Settings were selected: Compartments to Segment – nuclei; Edge Rules – discard cell if touching an edge; Nuclei-General Approach – object-based; Clean-up – fill holes; Signal Scaling – auto scale; Components for Nuclear Segmentation – hematoxylin, minimum signal = 0.37; Size Range – in pixels, minimum = 100, maximum = 10,000. In the IHC Settings tab, for Ki-67, Component was set to Purple, the Threshold Max was set to 1.0/Auto, and the Optical Density Positivity Threshold was set to 0.040. For Phh3, Component was set to Dab, the Threshold Max was set to 1.0/Auto, and the Optical Density Positivity Threshold was set to 0.30. Exported tables included Cell Segmentation Data and Score Data. Sectioning, immunohistochemistry, and imaging slides were completed at the University of Wisconsin Translational Research Services in Pathology core.

The metric used to quantify differences in cell division was the average inForm/Nuance percent-positive scores for each sample (averaged across 8-10 images). Differences in positivity between strains were determined using t-tests.

### Respiratory exchange ratio, food/water consumption, and physical activity

Seven to nine mice of each strain and sex were individually housed in a TSE Systems PhenoMaster metabolic cage from day 28 to 35. Data from the first and last days were dropped. Mice had *ad libitum* access to water and food. From day 21-28, mice were acclimatized in a metabolic cage fitted with PhenoMaster drinking and feeding modules.

Indirect calorimetry sampling was drawn from two experimental cages (with individually housed mice) and one control (empty) cage. Per cage, calorimetric sampling occurred every 9 minutes. Air flow rate was set to 0.22 liters/minute.

TSE Systems LabMaster software enabled continuous monitoring of O_2_ consumption and CO_2_ production. This permitted acquisition of the respiratory exchange ratio (RER), which is the ratio of metabolically produced CO_2_ exhaled to O_2_ consumed in a closed system compared to the same ratio in ambient (control) air. An RER near 0.7 indicates an organism is utilizing fats as a primary fuel source, whereas an RER near 1 indicates carbohydrates are the primary fuel source (Simonson & DeFronzo, 1990; Speakman, 2013). RER values in between these indicators signify that both fats and carbohydrates are utilized.

RER is an accurate reflection of the fuel used only if the mouse is at rest or engaging in steady-state, non-strenuous, aerobic exercise (Simonson & DeFronzo, 1990; Seidenberg & Beutler, 2008; Speakman, 2013). During strenuous anaerobic exercise, additional CO_2_ is exhaled to compensate for the increasing blood acidity brought on by lactic acid production (Seidenberg and Beutler, 2008). TSE Systems LabMaster software also recorded mouse activity behaviors. We defined a spectrum of data sets from rest to steady-state exercise to strenuous exercise. Activity was measured by quantifying non-wheel, ambulatory movements along the cage’s long axis, with a count defined as the interruption of two different light beams separated by 15 mm. Considering activity across the six days, we treated the average activity per hour for a mouse as a data point, pooled strain-specific mouse averages across hours, and compared strains with separate t-tests for each hour. We evaluated p-values using a range of critical values that correct for familywise error rates (Bonferroni, Holm, and Benjamini-Hochberg).

We predicted that if a dynamic pattern of RER reflects, either as cause or consequence, the differences in weight gain from days 29-34, then such a pattern would be most apparent in the restful/steady-exercise data sets and would be lost in those that reflect the most strenuous activities, where excess CO_2_ exhalation confounds RER interpretation. The rest-activity spectrum was organized into five classes. The first class is presented in the main text and remaining classes are presented in Supplementary Figures 4 and 5:

1. Circadian rest: with reference to the 24-hour clock, RER recorded during the four hours when all non-wheel activity and wheel-use, considered jointly, were at their lowest.
2. Rest-No activity: the collection of all RER records across the 24-hour clock for which no activity was recorded in the same time stamp.
3. Non-wheel Activity: all RER records across the 24-hour clock that were collected in time stamps with ambulatory movement and no wheel activity.
4. Wheel-Long run: RER recorded within periods when a large proportion of a mouse’s time on the wheel was spent in a continuous run. This category was created using three records: SumTime, which is the total time a mouse is in the wheel; MaxLen, which is the longest individual run interval, where a run is defined (by the user) as the duration of movement that exceeds and then drops below 25 rpm; and SumRuns, which is the total number of run intervals. For each strain and sex, we calculated a quotient, MaxLen/SumTime, and retained all values above the median as they represented long stretches of continuous running. This metric, MaxLen/SumTime, was then plotted against SumRun, and RERs associated with MaxLen/SumTime values below the median SumRun were retained for analysis. These data represented RER during continuous, low-exertion running bouts during intervals when run starts and stops were minimized.
5. Wheel-Spurt: RERs recorded within periods when a large proportion of a mouse’s time on the wheel was spent in bouts of starting and stopping. For each strain and sex, this category was created by calculating a quotient, SumRun/SumTime, and retaining quotients above the median value. These data represented RER recorded during long intervals of wheel use composed of many run starts and stops, thus capturing unsteady, high-exertion behavior.

We treated the average RER within a rest-activity class for a mouse as a data point, pooled strain-specific mouse averages across days, and compared strains using ANOVA with day as a main effect or as part of an interaction term with strain. Pooled mouse-averages were further evaluated with separate t-tests for each day. The p-values were evaluated using a range of critical values that correct for familywise error rates (Bonferroni, Holm, and Benjamini-Hochberg).

Each metabolic cage was fitted with drinking and feeding modules that measured consumption to the nearest hundredth of a gram. Minimum and maximum consumption thresholds were set to 0.01 mL and 0.10 mL/minute for drink and 0.01 g and 0.10 g/minute for food. Pooled mouse-averages for food and water consumption were evaluated with separate t-tests for each day. The p-values were evaluated using a range of critical values that correct for familywise error rates (Bonferroni, Holm, and Benjamini-Hochberg).

All statistical analyses were conducted using R software (R version 4.3.1) (R Core Team 2023).

## Results

### Daily weights pinpoint periods of accelerated growth in Gough mice

Adult Gough Island mice are larger and heavier than mainland mice when raised in a common environment (Gray et al., 2015) (Figure 1A). To characterize divergence in growth trajectories between Gough mice and mainland mice with high resolution, we measured daily weights for the first 42 days of postnatal development. Gough female body weight is distinguishable from that of mainland females beginning on day 13 after birth (t(46) = 2.63, *P* < 0.01) (Figure 1B). By day 42 (the last day of record), Gough females weigh an average of 46% more than mainland females (17.60 g vs. 12.05 g) (Figure 1B). Gough male body weight is distinguishable from that of male mainland mice beginning on day 7 after birth (t(53) = 2.94, *P* < 0.001) (Figure 1C). By day 42, Gough males weigh an average of 54% more than their mainland counterparts (20.98 g vs. 13.60 g) (Figure 1C).

**Figure 1.**
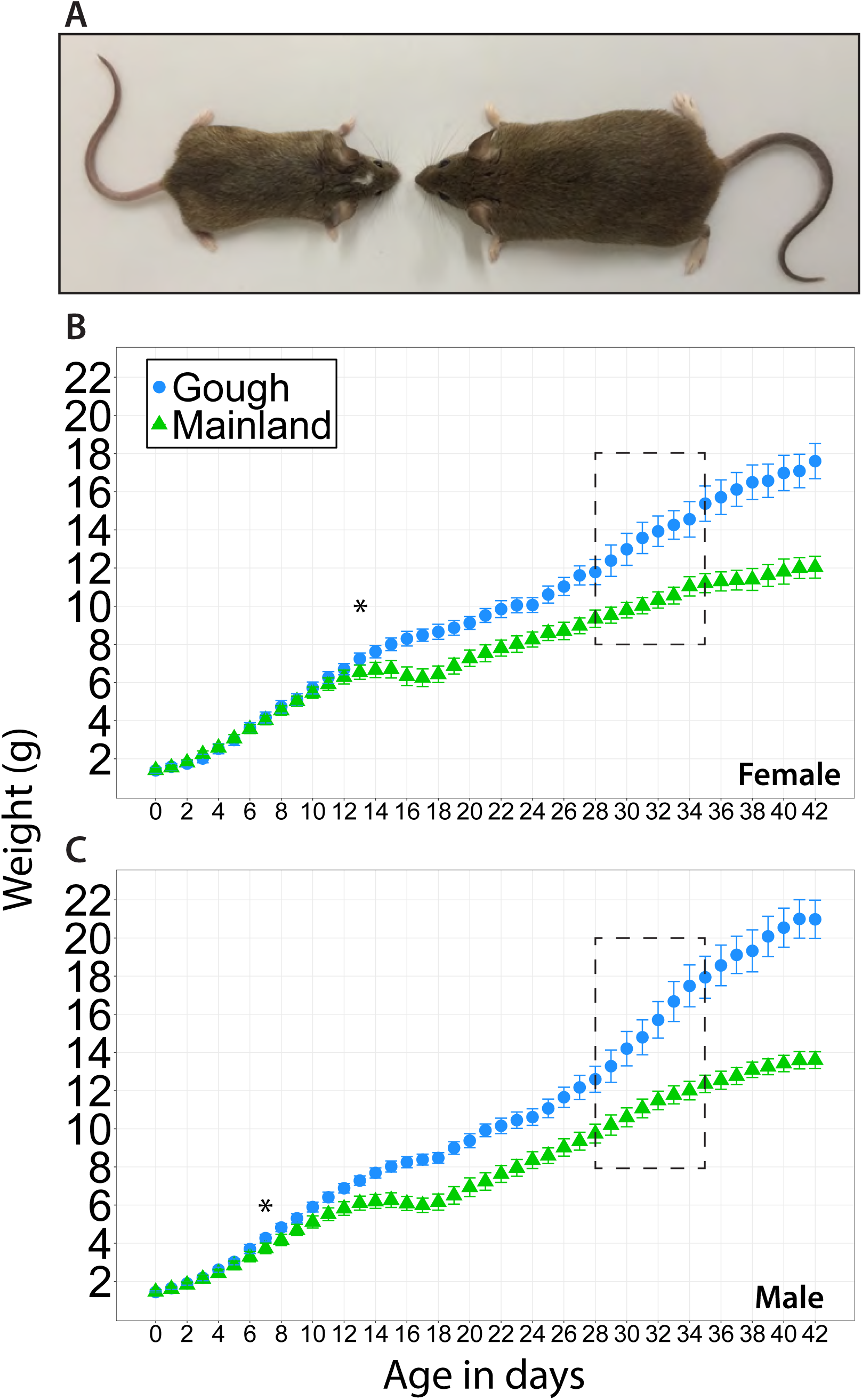
Weight differences in Gough Island and mainland house mice. (A) A sexually mature Gough Island mouse (right) photographed next to a mouse of the same age from a wild-derived inbred strain (WSB/EiJ). (B, C) Mean daily weight comparison. Color and shape designations are consistent across panels. Error bars are ± 2 x standard error of the means. Asterisks mark the first day of significant difference; all subsequent days exhibited significant differences between the strains. Dashed rectangles highlight week 5, a period of exceptional weight gain in Gough Island mice (see text).

From day 13 to day 18, mainland mice do not accumulate body weight, and even lose weight on days 15-17 (Figure 1B, C; Supplementary Figure S1). Growth of Gough mice is strikingly different during the same period, with mice continuing to accumulate weight (Figure 1B, C; Supplementary Figure S1). On day 13, female Gough mice weigh 11% more than mainland counterparts; by day 18, they weigh 35% more. On day 13, male Gough mice weigh 19% more than mainland counterparts; by day 18, they weigh 38% more (Figure 1B, C; Supplementary Figure S1A, B).

Widely divergent weight gain is observed again in the fifth week. Remarkably, during the fifth week, Gough females acquire more than 0.5 g in four separate 24-hour periods, whereas during the same interval mainland females do not acquire 0.5 g during any 24-hour period (Figure 1B; Supplementary Figure S1A). Similarly, Gough males gain more than 0.8 g in four separate 24-hour periods; mainland males never accumulate more than 0.47 g in a 24-hour period (Figure 1C; Supplementary Figure S1B).

### Organ weights and growth rates suggest key contributions of fat to body weight gain in Gough mice

We asked whether organ weights could shed light on the earliest instances of strain divergence in body weight. In both females and males, three visceral fat depots – gonadal, mesenteric, and retroperitoneal – are significantly heavier in Gough mice than mainland mice by 2 weeks, the earliest recorded period (Figure 2A, B; Supplementary Table S1). Gough mice also have heavier inguinal subcutaneous fat depots and inter-scapular brown fat depots than mainland mice at either 2 weeks or 3 weeks (Figure 2A; Supplementary Table S1). In addition, the intestine is longer in Gough mice than mainland mice by 2 weeks (Figure S2; Supplementary Table S1).

**Figure 2.**
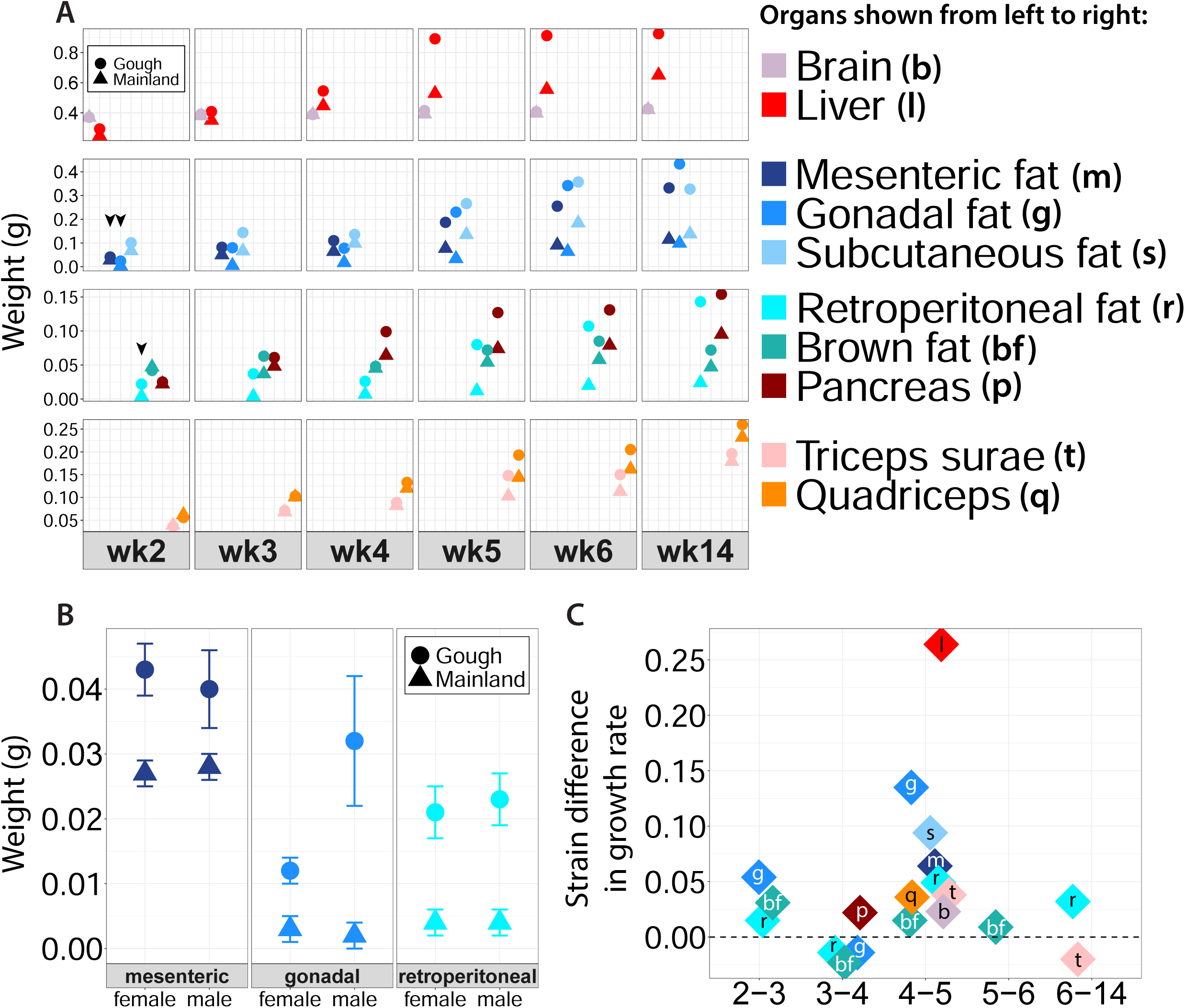
Weekly organ growth. Organ colors and abbreviations are consistent across panels A-C. (A) Sex-averaged weekly organ weights. Black arrowheads indicate weights highlighted in panel B. (B) Mean weight differences of 2-week visceral fat depots. All sex-specific comparisons are significantly different. Error bars are ± 2 x standard error of the means. (C) Significant differences between Gough mice and mainland mice in the rate of organ weight gain (differences in the slopes) for each weekly transition. Points above the dotted zero-line indicate weekly transitions when organ weight accumulation is greater in Gough mice than mainland mice. Points below the dotted zero-line indicate transitions when organ weight accumulation is greater in mainland mice than Gough mice.

Non-fat organs likely do not contribute as much as fat to early body weight differences between the strains. In males, the liver and pancreas are not significantly heavier in Gough mice until the third week; in females, these organs are not consistently different until 4 weeks and beyond (Figure 2A; Supplementary Table S1). Muscles (represented by the quadriceps and triceps surae) and the brain are heavier in Gough mice than mainland mice by 5 weeks (Figure 2A; Supplementary Table S1), a difference that is not consistently maintained (Figure 2A; Supplementary Table S1).

While the liver at 5 weeks yields the single largest strain difference in organ weights across assayed time points – with a 0.37 g excess in Gough mice compared to mainland mice – the relative contributions of visceral and subcutaneous fat depots compared to other organs expand over time (Figure 2A). For example, sex-averaged gonadal fat depots are 0.06 g, 0.20 g, 0.28 g, and 0.33 g heavier in Gough mice than mainland mice at 4, 5, 6, and 14 weeks, respectively (Figure 2A).

When considering differences in weekly organ growth rates between Gough mice and mainland mice, three salient patterns emerge (Figure 2C; Supplementary Table S2). First, across all organs except the pancreas, growth rate is higher in Gough mice than mainland mice from 4 to 5 weeks, indicating that this seven-day period is a key temporal window for the manifestation of the island rule in Gough mice (Figures 2C; Supplementary Table S2). Second, the brown, gonadal, and retroperitoneal fat depots grow faster in Gough mice than mainland mice from 2 to 3 weeks. This pattern suggests that visceral and brown fat depots are key contributors to the elevated body weight in Gough mice relative to mainland mice during the third week (Figure 1; Figure 2A, C; Supplementary Table S2). Third, from week 6 to 14 the retroperitoneal fat depot is the only organ with a greater growth rate in Gough mice (Figure 2C; Supplementary Table S2), indicating a special role for this fat depot in furthering the divergence in body weight between Gough mice and mainland mice into adulthood.

The effect of mouse strain on the growth of most organs does not differ between the sexes (Supplementary Table S3). Exceptions are brown, gonadal, and retroperitoneal fat, and intestine length (Supplementary Tables S3; Supplementary Table S4).

### Blood chemistry suggests metabolic factors tied to increased body weight in Gough mice

To investigate endocrinological factors associated with differential growth between Gough mice and mainland mice, we examined eight plasma analytes with established connections to body size. Four of the eight analytes we quantified are significantly different between strains at 1 week of age (Table 1). Sex-averaged leptin, free fatty acids, and Igf-1 concentrations are lower in Gough mice than mainland mice, while ghrelin is higher. These strain differences precede or co-occur with the earliest strain differences in body weight and organ weights (Figure 1; Figure 2).

**Table 1.** Effects of strain on plasma analytes at 1 week of age (sexes combined). T-test.

|  |  |  |  |  |
| --- | --- | --- | --- | --- |
| Plasma analyte<sup>a</sup> | Mean (Standard deviation) |  | t<sup>f</sup> | p-value* |
|  | Mainland | Gough |  |  |
| Ghrelin<sup>b</sup> | ***781.2 (172.8)*** | ***1752.4 (203.3)*** | ***3.5 (16)*** | ***< 0.01*** |
| Leptin<sup>b</sup> | ***586.8 (127.7)*** | ***90.7 (16.5)*** | ***3.9 (18)*** | ***< 0.01*** |
| Glucose<sup>c</sup> | 20.7 (1.2) | 18.8 (2.3) | 0.7 (17) | 0.477 |
| Adiponectin<sup>d</sup> | 2090.3 (315.8) | 1583 (176.9) | 1.5 (16) | 0.159 |
| IGF-1<sup>d</sup> | ***28.4 (2.9)*** | ***14.1 (2.3)*** | ***3.8 (16)*** | ***< 0.01*** |
| Free fatty acid<sup>e</sup> | ***0.13 (0.01)*** | ***0.08 (0.01)*** | ***4.6 (15)*** | ***< 0.001*** |

<sup>a</sup> Concentrations for Glucagon and Insulin were below the detection threshold.
<sup>b</sup> pg/ml; <sup>c</sup> mg/dl; <sup>d</sup> ng/ml; <sup>e</sup> nM.
<sup>f</sup> Degrees of freedom in parentheses.
\* Significant results are bolded and italicized.

As preliminary ANOVA indicated that differences in analyte concentrations between Gough mice and mainland mice depend on sex after 1 week of age, analytes were evaluated separately in females and males at all ages beyond 1 week (Supplementary Table S5). These analyses reveal four groups of plasma analytes that show distinct temporal patterns (Figure 3; Supplementary Table S5). First, in both sexes, levels of ghrelin, leptin, glucagon, and glucose are higher in Gough mice than mainland mice when there are significant differences between strains (Figure 3A). Given that ghrelin promotes hunger, while leptin promotes satiety (Uchida et al., 2013; Cui et al., 2017), we asked whether the ratio of these analytes differs between strains (Figure 3B). When considering this ratio, two signals shared across the sexes stand out: a higher ghrelin to leptin ratio in Gough mice at 2 weeks and the lack of strain differences in this ratio during key periods of body growth and organ growth that span weeks 4 and 5 (Figure 1; Figure 2; Figure 3B; Supplementary Figure S1).

**Figure 3.**
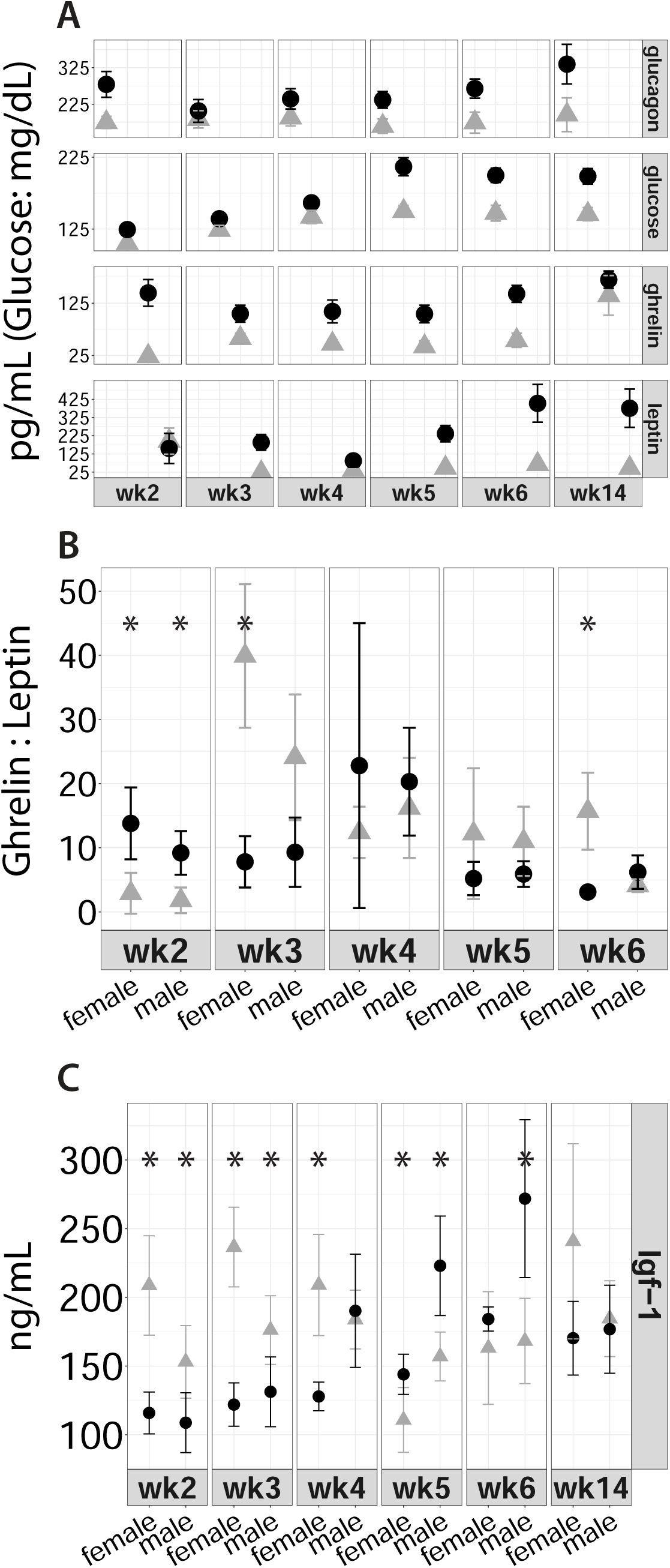
Weekly fasting plasma analyte concentrations. Black circles represent Gough Island mice; gray triangles represent Mainland mice. (A) Sex-averaged plasma analyte concentrations. Ghrelin and leptin concentrations are reduced by a factor of 100 and 10, respectively. Only week 3-glucagon and week 2-leptin concentrations are not significantly different in either females or males. (B) Mean weekly plasma ghrelin:leptin ratios. Week 14 data not shown because large variance distorted pattern observed across weeks 2-6; qualitatively, week 14 is similar to week 6 ratios. (C) Mean weekly Igf-1 concentrations. In all panels, error bars are ± 2 x standard error of the means and, when shown, asterisks mark significant differences.

In the second group, adiponectin and Igf-1 exhibit complex temporal sex-specific and strain-specific plasma concentrations (Supplementary Table S5). Soon after birth, both analytes are lower in Gough mice than mainland mice (Figure 3C; Supplementary Figure S3). Then, beginning at 4 weeks (adiponectin) or 5 weeks (Igf-1), and beyond, there are periods when concentrations in Gough mice are significantly higher than those of mainland mice (Figure 3C; Supplementary Figure S3). The higher concentrations of adiponectin at later time points parallel the increasing weight of fat depots (Figure 2; Supplementary Figure S3). Given that Igf-1 promotes growth (Stratikopoulos et al., 2008; Lui and Baron, 2011), our results do not support a major role for this analyte in the earliest differential weight gain between Gough mice and mainland mice (Figure 1; Figure 3C; Supplementary Figure S1). In contrast, higher Igf-1 levels in Gough mice than mainland mice beginning at week 5 suggests a potential role for Igf-1 during this astounding period of differential weight gain (Figure 1; Figure 2; Figure 3C; Supplementary Figure S1).

The remaining two salient patterns of blood chemistry come from single analytes. Gough mice and mainland mice do not differ in insulin concentration from 2 to 6 weeks, nor at 14 weeks (Supplementary Table S5). Furthermore, among females, Gough mice have lower levels of free fatty acids than mainland mice across ages, whereas there is no strain difference in males (Supplementary Table S5).

### Rates of cell division illuminate cellular mechanisms of organ-specific weight gain in Gough mice

To determine whether elevated rates of cell division contribute to larger body size in Gough mice, we applied immunohistochemistry to the liver and the gonadal fat depot at 15 days in both sexes, and again at 30 days in females and 32 days in males. On day 15, female and male Gough livers exhibit significantly more cell division than mainland livers when evaluated with Ki-67 (Figure 4), which labels cycling cells. Higher cell division on day 15 is further indicated in female Gough livers using Phh3 (Figure 4), which also labels cycling cells. Additionally, on day 32, Ki-67 labeling in Gough male livers shows greater cell division than in mainland livers (Figure 4). Though it does not reach statistical significance, this pattern is mirrored with Phh3 (Figure 4).

**Figure 4.**
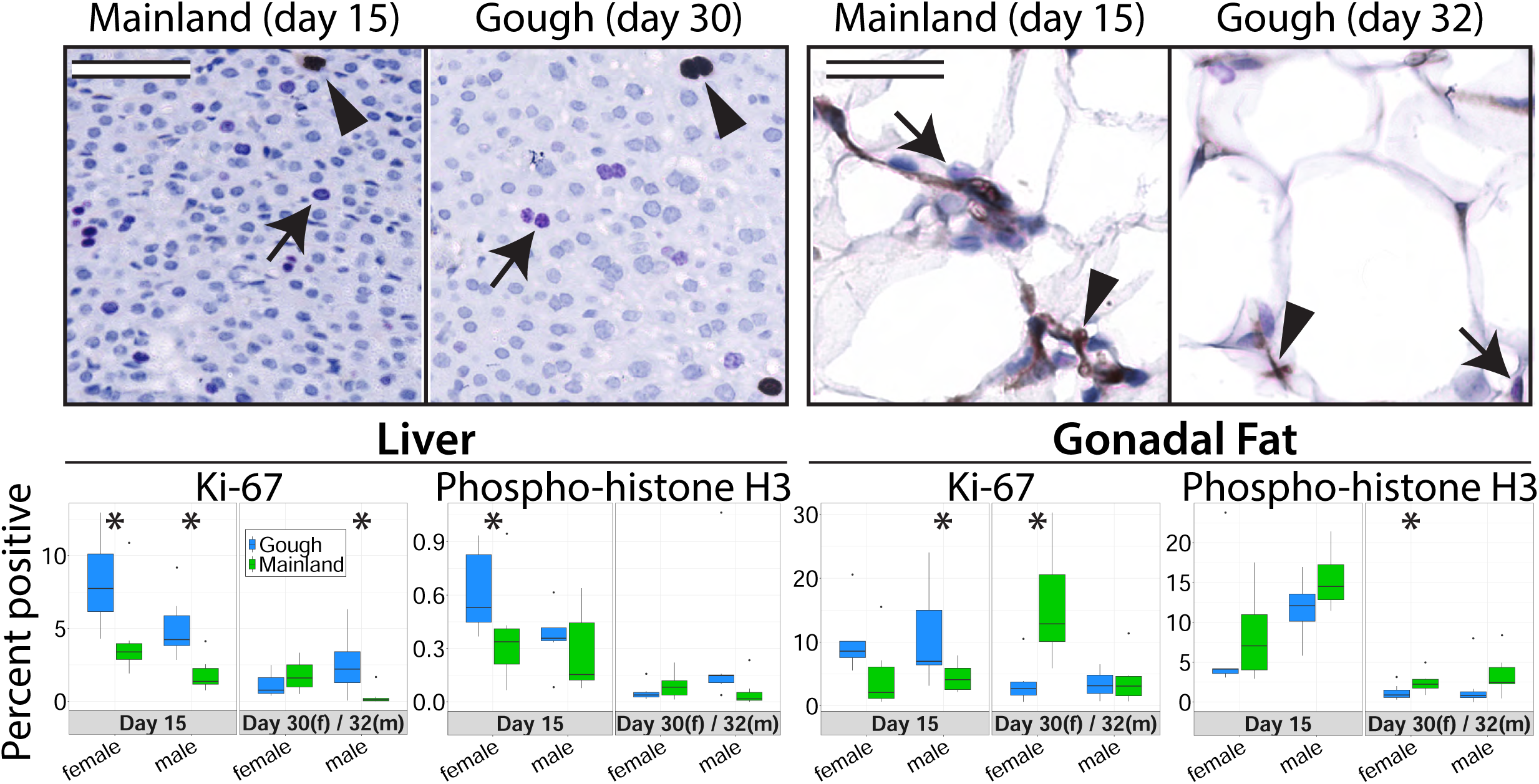
Immunohistochemistry (IHC) images and quantification for Ki-67 and Phh3. Images show example dual-staining of Ki-67 and Phh3 in liver and gonadal fat. All images for the liver are from females. All images for the gonadal fat are from males. Liver scale bar is 50 um. Gonadal fat scale bar is 25 um. Arrows indicated Ki-67-positive cells in all IHC images, while arrowheads indicated Phh3-postive cells in all IHC images. In box plots, asterisks indicate p < 0.1. Color designations are consistent across box plots; in all comparative sex-by-time point box plots, Gough data is on the left and Mainland data is on the right.

Patterns of cell division are more complicated in 15-day gonadal fat depots. On day 15, Ki-67 positivity indicates that male Gough mice experience more cell division than mainland male mice (t(11) = 2.1, *P* < 0.1; Figure 4). A trend in this direction is also visible in females (t(9) = 1.6, *P* = 0.1). Phh3 suggests the opposite pattern in 15-day male gonadal fat (Figure 4), though the strain difference does not reach significance (t(11) = 1.8; *P* = 0.1). One useful perspective to consider when comparing these results is that while Ki-67 and Phh3 both label cycling cells, Ki-67 labels the longer G1, S, and early G2 phases, whereas Phh3 labels the shorter late G2 and mitotic phases (Hendzel et al., 1997; Van Hooser et al., 1998; Maccallum et al., 2000; Crosio et al., 2002; Li & Spalding, 2022). This distinction may have implications for differential growth of the gonadal fat depot in Gough mice and mainland mice, with more Gough adipocytes spending time in the longer phases of the cell cycle (labeled by Ki-67), and mainland adipocytes moving through the cell cycle at a relatively faster pace (indicated by elevated M-phase-labeling with Phh3). At 30 days, female Gough gonadal fat shows lower cell division than female mainland gonadal fat (Figure 4). During the 24-hour period from day 29 to 30, Gough females add significantly more weight than mainland females (Figure 1B; Supplementary Figure S1). One cellular mechanism by which fat depots expand in size is by exiting the cell cycle and upregulating lipid production and storage (Li & Spalding, 2022; Sakers et al., 2022).

### Gough mice increase reliance on carbohydrates and reduce reliance on fat for fuel

To evaluate the use of carbohydrates and fats as fuel sources, we used indirect calorimetry. We focused on the fifth week of postnatal growth because of exceptional growth differences during this period (Figure 1B, C; Supplementary Figure S1). From day 29 to day 34, Gough mice exhibit climbing average pooled Respiratory Rate Ratios (RER), indicating a shift towards carbohydrates as a preferred fuel source (Figure 5). This trend is especially visible for the restful activity classes (Figure 5; Supplementary Figure S4; Supplementary Figure S5; Supplementary Table S6). For example, in the Circadian Rest class, the average day-29 RER is 0.80 and 0.79 for Gough females and males, respectively, and by day 34, the average RER is 0.88 and 0.89 for females and males, respectively (Figure 5Ai, Bi; Supplementary Table S6). In contrast, average RER in female and male mainland mice either decreases from day 29 to day 34 or remains indistinguishable from day to day (Figure 5Ai, Bi; Supplementary Table S5). Additionally, for the Circadian Rest (both sexes) and Rest-No Activity (females) classes, there is no significant strain difference in RER on day 29, but by day 34, Gough mice exhibit significantly higher RER than mainland mice (Figure 5Ai, Bi; Supplementary Table S6). This pattern is echoed in sex-specific ANOVAs that assess the effects of strain, day and their interaction (Supplementary Table S7): the effect of day on RER is different between Gough mice and mainland mice.

**Figure 5.**
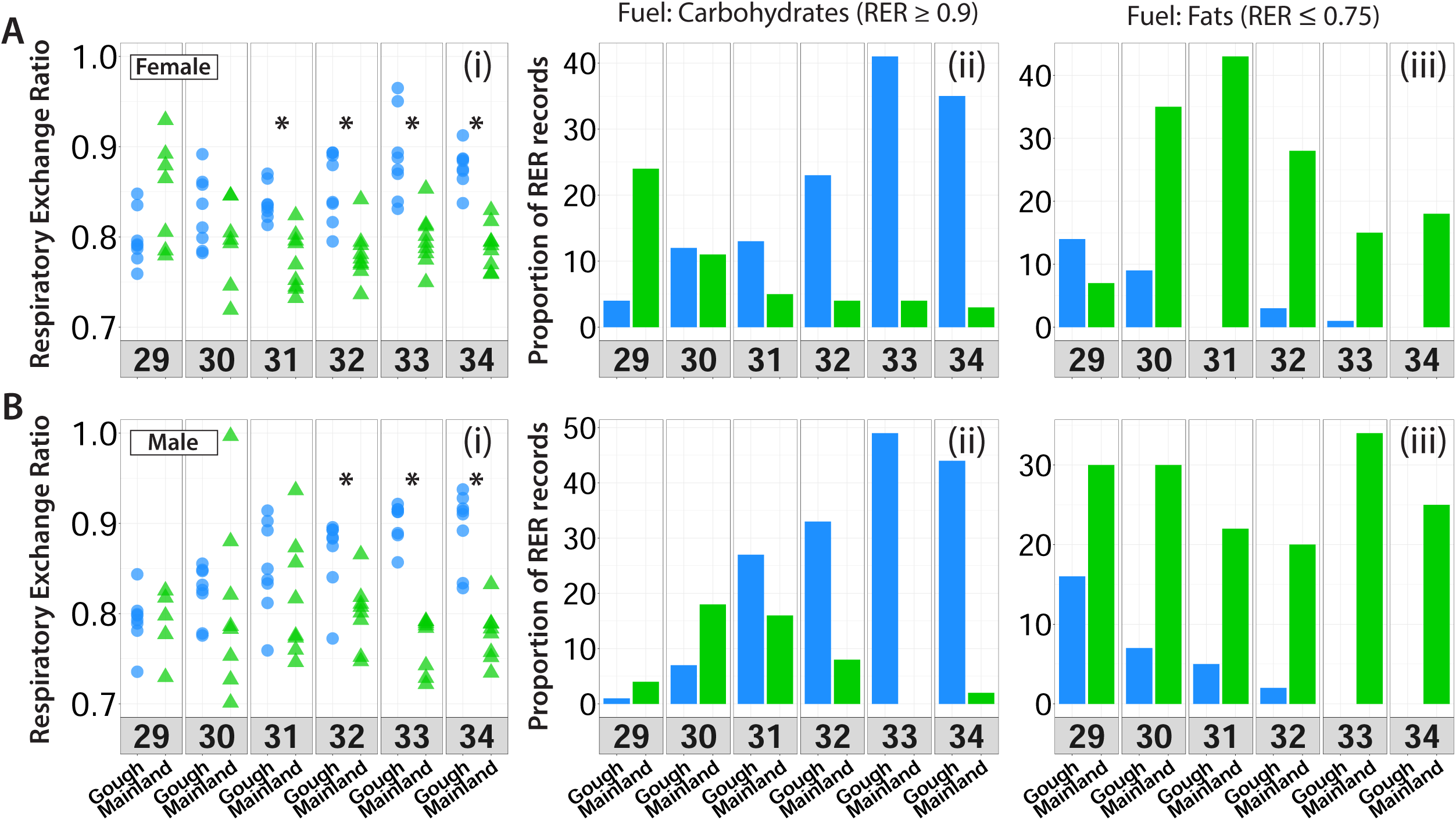
Comparison of Circadian Rest respiratory exchange ratios (RER; see Methods) across post-natal days 29-34. The color blue and circles represent Gough Island mice; the color green and triangles represent Mainland mice. (Ai-iii) Females. (Bi-iii) Males. (Ai, Bi) Scatter plots treat the mean RER for a mouse as a data point and pool strain-specific mouse means for each day. Asterisks mark significant differences. (Aii, Bii) The proportion of RERs that are categorically indicative of energy production via carbohydrates. (Aiii, Biii) The proportion of RERs that are categorically indicative of energy production via fats.

To better characterize the pattern of a climbing daily RER in Gough mice, we visualized the proportion of RER records for each strain, aggregated from all mice in that strain, that are categorically indicative of exclusive carbohydrate use (RER ≥ 0.9) or exclusive fat use (RER ≤ 0.75) for energy production (Figure 5). Upon partitioning the daily RER records in this manner, a salient pattern emerges that may be a cause or consequence of the strain differences in weight gain. For Gough mice, in the Circadian Rest class, the proportion of RER values that indicate carbohydrate use rises from day 29 to day 34 (Figure 5Aii, Bii). Concurrently, RER records of carbohydrate use in mainland mice either decrease or remain steady (Figure 5Aii, Bii). For example, in female and male Gough mice the proportion of day 29 RER records ≥ 0.9 is 4% and 1%, respectively; by day 34 the proportion is 35% and 44%, respectively (Figure 5Aii, Bii). In contrast, for mainland mice, on day 29, the proportion is 24% and 4% for females and males, respectively; on day 34, the proportion is 3% for females and 2% for males (Figure 5Aii, Bii). The patterns of carbohydrate fuel use observed in the Circadian Rest class are reflected and confirmed in additional rest-activity classes (Supplementary Figure S4; Supplementary Figure S5).

When considering fat metabolism, a different and complementary pattern emerges. For Gough mice, in the Circadian Rest class, the proportion of RER records indicative of fat fuel use (RER ≤ 0.75) diminish rapidly and completely from day 29 to 34; for female and male Gough mice there are zero RER records ≤ 0.75 on day 34 (Figure 5Aiii, Biii). In contrast, female mainland mice exhibit successive days of diminishing fat fuel use (days 31-33, for example) but not complete abandonment of this fuel source (Figure 5Aiii). Fat fuel use in mainland males also contrasts with the pattern presented in Gough mice. In the Circadian Rest class, RER records from mainland male mice exhibit a more static pattern of fat fuel use from day 29-34: the proportion of RER records ≤ 0.75 across days 29-34 does not dip below 20% (Figure 5Biii). The patterns of fat fuel use observed in the Circadian Rest class are reflected and confirmed in additional rest-activity classes (Supplementary Figure S4; Supplementary Figure S5). In summary, in the fifth postnatal week, relative to mainland mice, Gough mice display an increasing reliance on carbohydrates as energetic fuel, while simultaneously reducing their dependency on fat.

### Gough mice are more active

To gauge activity, we measured ambulatory movement. Gough mice exhibit more ambulatory movement than mainland mice in 9 hours (females) or 10 hours (males) of the 24-hour clock (Figure 6). Mainland female and male mice are more mobile during only 2 of the hours. Considering total activity across all evaluated days for a mouse as a data point and pooling data points by strain confirms that Gough mice are more active than mainland mice across days 29-34 (female: t(12.7) = 6.8, *P* < 0.001; male: t(8.8) = 4.5, *P* < 0.001). Our results show that during a period in which Gough mice gain significantly more weight than their mainland counterparts, they are also significantly more active (Figure 1B, C; Figure 6; Supplementary Figure S1).

**Figure 6.**
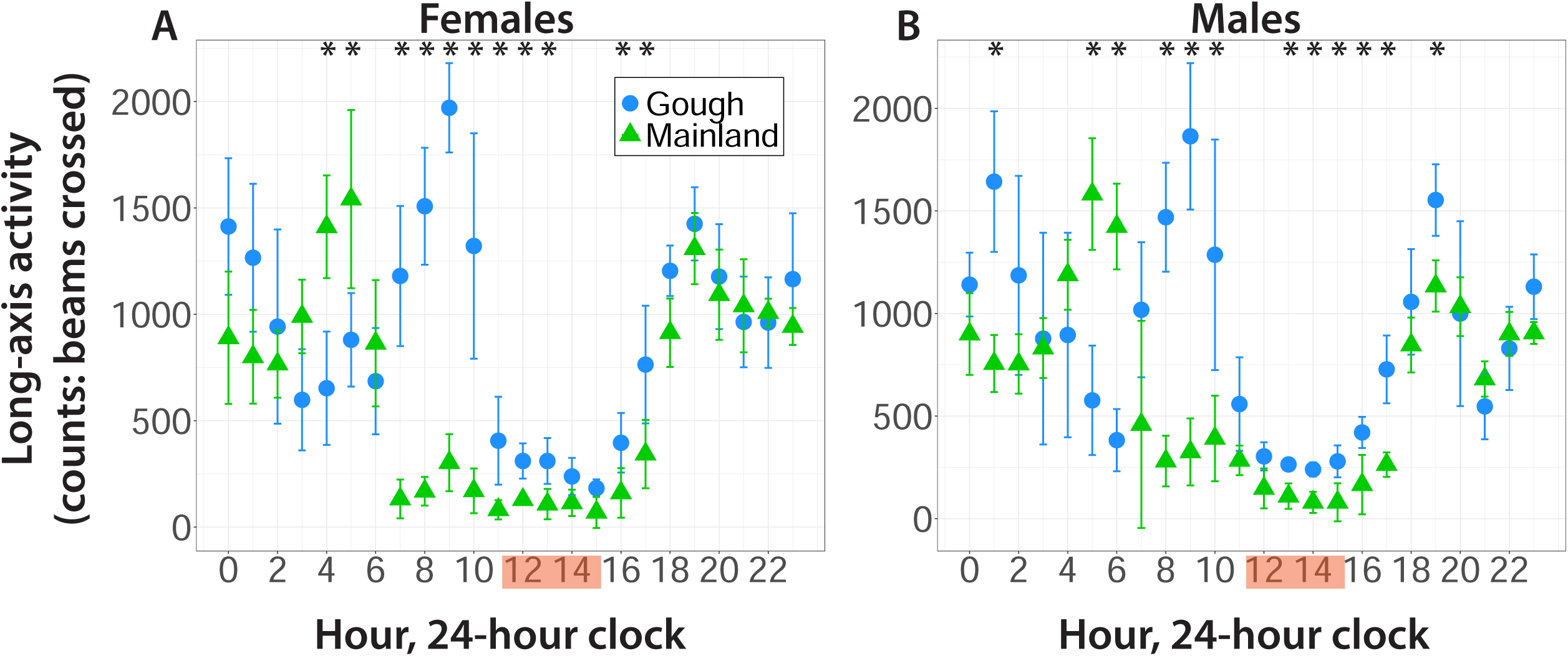
Activity patterns across days 29-34. (A, B) Long-axis activity is non-wheel ambulation/movement. Mean activity per hour per mouse across days 29-34 were pooled to obtain strain-specific means and variances across hours. Orange highlighted intervals mark the four hour interval used for the Circadian Rest rest-activity class in Figure 5. Asterisks mark significant differences and error bars are ± 2 x standard error of the means.

### Gough mice do not eat more food, though they drink more water

While male Gough mice consume less food than male mainland mice on days 29-31 (Supplementary Table S8), the total amount of food consumed across days 29-34 does not differ between males from the two strains (t(10) = 1.8, *P* = 0.1). Gough females and mainland females show no statistical difference in daily (Supplementary Table S8) or cumulative food consumption (t(9) = 0.6, *P* = 0.6). In contrast, Gough mice drink more water on both a daily basis (Supplementary Table S8) and a cumulative basis (female: t(7.4) = 6.3, *P* < 0.001; male: t(6.2) = 5.2, *P* < 0.01).

## Discussion

### Contrasting avenues of body weight evolution on remote islands and the continental mainland

Organismal body weight mediates and reflects interactions with the environment. Empirical and theoretical studies suggest that body weight impacts reproductive success, directly linking it to fitness and natural selection (Young, 1976; Lindstedt & Boyce 1985; Blanckenhorn, 2000; Davidowitz et al., 2005; Bernstein, 2010; Little, 2020; Bright Ross et al., 2021). Body weight is connected to diet, resource proximity, temperature exposure, intraspecific and interspecific competition, and predation risk, among other environmental variables (Blanckenhorn, 2000; Bernstein, 2010; Bright Ross et al., 2021).

Gough Island is one of the remotest land masses on the planet and its unique environment likely played an important role in the evolution of extreme body weight in mice on the island. Gough island is positioned more than 2,400 kilometers from mainland South Africa and South America, is home to no predators of house mice (Cuthbert et al., 2016), is void of manmade structures that could serve as commensal shelters for house mice (with the exception of a small meteorological station) (Rowe-Rowe & Crafford, 1992; Jones et al., 2003), and is host to a unique assemblage of food resources, including grasses, ferns, lumbricid worms, moths, and seasonally fluctuating access to seabird chicks (Jones et al., 2003; Cuthbert & Hilton, 2004; Cuthbert et al., 2016).

The unusual environment of Gough Island could generate selection for new behaviors. Indeed, behavioral assays showed that Gough mice are more willing to explore a novel environment than mainland mice (Stratton et al., 2021; Stratton et al., 2023). Remarkably, Gough mice seem to have evolved large bodies without the specific behavioral shifts one might predict: eating more and moving less. Compared to mainland mice, Gough mice are more active and eat the same amount of food during the 5^th^ week of age. Given that Gough mice are already heavier than mainland mice from week 4-5, their increased relative activity without elevated food consumption might be expected to incur energy costs that would reduce growth and body size maintenance. In fact, the opposite is true. Therefore, Gough mice activity levels and feeding behavior are potentially uncoupled from metabolic mechanisms that promote weight accumulation.

Respiration measurements indicate that compared to mainland mice, Gough mice rely more on pure carbohydrate metabolism and less on pure fat metabolism for energy production. Hence, one evolved mechanism of weight gain in Gough mice is the attenuation of fat catabolism as an energy source. Isolated islands away from continental fauna might favor more active animals that use opportunities to forage for food and expand territories made available by the absence of predators and interspecific competitors (Charnov & Berrigan, 1993; Bernstein, 2010). But an active animal would experience higher caloric expenditure, presumably promoting leanness (Bernstein, 2010). To counter an active caloric expenditure and maintain greater body weight, island populations would need to evolve metabolisms that favor burning fuel sources other than fat, such as the shift towards carbohydrate usage in Gough mice. More generally, isolated islands without predators or interspecific competitors may favor larger rodents with altered metabolic pathways in organs that create, store, and process fats (such as the liver and fat depots). Genetic variants associated with fat metabolism and body size have been identified in relation to size differences among mainland populations of house mice, including gene regulatory divergence in the liver (Mack et al., 2018; Phifer-Rixey et al., 2018; Ferris et al., 2021; Ballinger et al., 2023; Durkin et al., 2024; Gutiérrez-Guerrero et al., 2024). Moreover, several genes implicated in body size differences among mainland house mice (Mack et al., 2018; Ballinger et al. 2023; Gutiérrez-Guerrero et al., 2024) are also differentially expressed in the livers of Gough mice and mainland mice (Nolte et al., 2020). These genes include *Baz2a, B4galnt1, Cdk1, Ctdsp2, F11, Inhbe, Itga1, Mroh7, Myl6, Pid1, Scar1b, Slc35d1, Smarcc2, Snx25, Timeless, Tsfm,* and *Vdac1*.

Isolated islands are also less likely to offer human-commensal structures for nesting and rest, a major contrast from mainland environments for house mice. Adaptations for constructing or seeking shelter in less-protected environments may favor large body size, with ample fat depots to ward off the cold. Inbred house mouse strains derived from North American populations at high latitudes weigh more than inbred strains from lower latitudes, reflecting adaptation to cooler temperatures and following Bergmann’s rule (Lynch, 1992; Bernstein, 2010; Phifer-Rixey et al., 2018; Ferris et al., 2021; Ballinger & Nachman, 2022). Given that Bergmann’s rule extends to mainland house mice with access to human-commensal structures that mitigate the impact of cold temperatures, similar ecological drivers likely contribute to body weight evolution in rodents from remote islands devoid of human inhabitants. The higher growth rate of the brown scapular adipose depot in Gough mice could constitute an adaptation to the cold (Cannon & Nedergaard, 2004; Van Sant & Hammond, 2008; Velotta et al., 2016; Ballinger & Andrews 2018). In laboratory colonies, house mouse pups are independently mobile and often weaned at 3 weeks. To the extent that this habit reflects natural dispersal behavior, the larger brown adipose depot at 3 weeks might buffer young Gough mice against a more exposed environment and contribute to their overall greater body weight. Gene regulatory evolution documented in the brown fat of adult house mice has been implicated in climatic adaptation, including body size differences among mainland populations of house mice (Ballinger et al., 2023; Durkin et al., 2024). Our results suggest that brown fat growth and metabolism are targets for gene regulatory evolution in prepubescent island-adapted house mice.

Mice and other small rodents on islands with similar resource distributions, levels of predation, and shelter access could evolve greater body weight in similar ways to Gough mice. Phenotypic and developmental dissection of other populations with exceptional body weights would be a productive path toward revealing general mechanisms responsible for the island rule.

### Endophenotypic basis of body weight evolution

Organismal body weight is a composite phenotype in two basic ways. First, body weight is the sum of the weights of an organism’s constituent organs. Second, body weight at any given age is an ontogenetic trait, reflecting the accumulation of developmental and physiological processes through time (Atchley et al., 2000; Cheverud, 2005; Kenney-Hunt et al., 2006; Little, 2020). Therefore, understanding evolutionary transitions in body weight ultimately requires dissection of its sub-organismal components in an ontogenetic context.

Components of body weight that underpin the evolution of this trait are usefully viewed as “endophenotypes” or “intermediate phenotypes.” Originally applied to the effects of chromosomal polymorphism on meiosis in Drosophila (John & Lewis 1966), the endophenotype concept has provided insights into human disease (Gottesman & Gould 2003; Gould & Gottesman 2006; Bihoreau et al., 2017), animal sociality (Toth & Robinson, 2007; Rittschof & Robinson, 2016), and evolutionarily divergent traits (Landry & Rifkin, 2012; Lema & Kitano, 2013). The capacity for endophenotypes to connect a downstream phenotype of interest (e.g. body weight) with its genetic causes (Landry & Rifkin, 2012; Rittschof & Robinson, 2016) provides a strong motivation for identifying these characteristics.

Previous research discovered multiple quantitative trait loci (QTL) involved in the evolution of extreme body weight and related skeletal expansion in Gough mice, with genetic effects that vary across development and concentrate during the earlier weeks of the postnatal growth trajectory (Gray et al., 2015; Parmenter et al., 2016; Payseur et al., 2023). This dynamic genetic architecture suggests that body weight evolution involved changes to multiple components of body weight (see also Cheverud, 2005; Kenney-Hunt et al., 2006). By documenting the ontogenetic contributions of organ growth, endocrinological cues, physiological processes, and behavior to divergent weight accumulation in Gough mice and mainland mice, we discovered evolutionary endophenotypes replete with anatomical and temporal specificity. These endophenotypes could be regulated by distinct genetic mechanisms and acted upon independently by evolutionary forces. For example, distinct temporal patterns of divergence in fat and in circulating hormone levels suggest that these endophenotypes provide complementary insights into the evolution of extreme body size.

In Figure 7 and for the remainder of the Discussion, we use this endophenotypic perspective to propose a heuristic model describing how Gough mice evolved the gigantism intrinsic to the island rule. Our results provide evidence for heterochronic shifts in fat accumulation. With the exception of gonadal adipose depots, subcutaneous white, visceral white, and brown adipose depots begin differentiation during embryogenesis, though proportionally more maturation occurs postnatally (Han et al., 2011; Wang et al., 2013). That Gough mouse visceral fat depots (particularly gonadal adipose) are larger so early in postnatal development indicates a heterochronic shift. This shift is mediated by earlier developmental onset and perhaps a greater growth rate towards more rapid expansion and maturation of fat in Gough mice relative to mainland mice (in our study) as well as the classical strain C57BL/6J (Alberch et al., 1979; Han et al., 2011; Wang et al., 2013). The striking peramorphic (adult-like) nature of Gough visceral fat depot growth early in postnatal development is further emphasized by noting that among the 2-week cohort, one female mainland mouse had no retrievable gonadal fat and one mainland male had no retrievable gonadal or retroperitoneal fat. The evolved heterochronic changes in Gough adipose depots observed early in postnatal growth persist into puberty and adulthood. Fat accumulation in Gough mice is a consistent driver of their extreme weight throughout postnatal maturation.

**Figure 7.**
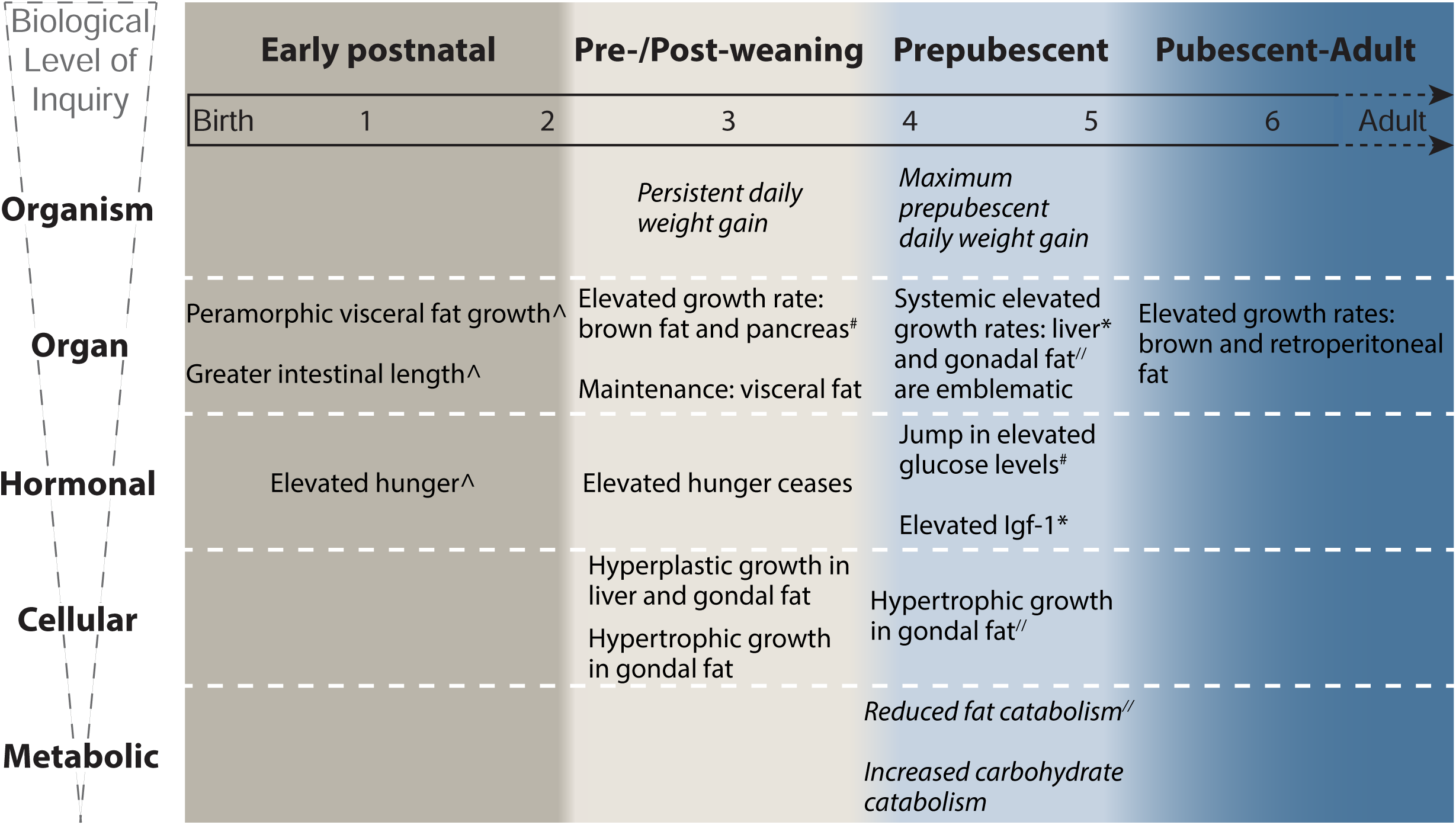
Genesis of Gough Island gigantism in four developmental phases. Phase boundaries were determined by transitions in and constellations of co-occurring endophenotypic measurements. Developmental and endophenotypic observations are provided from the perspective of a Gough Island mouse relative to a Mainland mouse. The timeline beginning with birth is shown in weeks, with dashed black lines indicating period between 6 and 14 weeks. Italicized text in grid indicates both a temporal resolution of 24 hours (daily resolution) and that documentation of the first instance of the trait’s divergence is reported in the main text. All other observations were inferred from week-to-week measurements and/or their first instance of divergence was not determined. Superscript symbols (^, #, *, //) link observations that may be biologically connected.

Our data reveal potential cellular and biochemical underpinnings of heterochronic alterations to adipose in Gough mice compared to mainland mice, including indications of hypertrophic adipocyte growth and a reduction in the use of fat as a fuel source. The salient empirical differences between Gough and mainland adipose depots implicate these organs as key evolutionary targets foundational to the morphological divergence of these populations. This finding resonates with recorded or predicted intraspecific differences in fat depot size and/or their molecular genetic regulation that are associated with differential survival (Lindstedt & Boyce, 1985; Harding et al., 2005; Bright Ross, et al. 2021) and environmental influences (Mastrangelo et al., 2019; Weldenegodguad et al., 2021; Xu et al., 2023).

Previously, Nolte et al. (2020) showed that the percentage of genes transcribed in the gonadal fat that were differentially expressed between Gough Island mice and mainland mice was greater than in the liver or hypothalamus, suggesting that gene regulatory evolution in fat depots plays an important role in body size evolution (Phifer-Rixey et al., 2018; Ballinger et al., 2023; Durkin et al., 2024), including the island rule (Nolte et al., 2020).

In comparison to adipose endophenotypes, endocrinological endophenotypes are more temporally punctuated. For example, higher ghrelin and ghrelin:leptin ratios at 1 and 2 weeks of age suggest that Gough pups might be hormonally predisposed to suckle and eat more than mainland pups (Uchida et al., 2013). As the stomach and duodenum secrete ghrelin, it is intriguing that Gough mice have longer intestines than mainland mice at 2 weeks, near to or coincident with their first divergence in body size (Korn, 1992; Date et al., 2000; Sykaras et al., 2014; Duque-Correa et al., 2021). Along these lines, follow-up studies that quantify differences in ghrelin secretion, pup nursing bouts, and milk content between Gough mice and mainland mice would be valuable. Maternal contributions to differences in pup weight go beyond milk content and can include effects from the uterine environment to postnatal mother-pup behavioral interactions. In our study, we did not explore maternal effects and cannot rule out maternal contributions to the phenotypic differences we detected. At the same time, previous studies of house mouse lines artificially selected for body size found that maternal effects were smaller than genetic effects and diminished with age (White et al., 1968; Moore et al., 1970).

Contrasting levels of ghrelin in Gough mice relative to mainland mice might also reflect evolutionary changes in other behavioral and metabolic processes independent of food intake. Ghrelin promotes locomotor activity akin to foraging behavior in rodents (Keen-Rhinehart & Bartness, 2005; Drazen et al., 2006; Jászberényi et al., 2006; Blum et al., 2009; Healy et al., 2011) and modulates migration-related decisions in birds (Goymanna, et al. 2017). Additionally, in mice, genetic manipulation of ghrelin or its signaling pathway disrupts reward-related and stress-related eating without strong effects on homeostatic feeding (Uchida et al., 2013; Cui et al., 2017). These behavioral roles of ghrelin are intriguing in light of discoveries that Gough mice predate on nesting sea birds (Cuthbert & Hilton, 2004; Cuthbert et al., 2016; Caravaggi et al., 2019) and exhibit augmented exploratory and mobility behavior (Stratton et al., 2021, 2023).

Another punctuated endocrinological endophenotype is the transition of higher plasma Igf-1 concentrations from mainland mice to Gough mice between the 4^th^ and 5^th^ week of age. This transition corresponds to another endophenotype: the near universal increase in organ growth rate in Gough mice during the same interval. This correspondence is consistent with an important, temporally restricted role for Igf-1 in postnatal growth in Gough mice after week 4 (Stratikopoulos et al., 2008; Lui & Baron, 2011). The evolutionary significance of Igf-1 genetic variants, plasma concentrations, and associations with life-history traits, including body size, has been investigated across a wide array of vertebrates (Dantzer & Swanson, 2012; Swanson & Dantzer, 2014; Lewin et al., 2017; Montoya et al., 2022; Plassais et al., 2022; Silva et al., 2023). Our findings are unique for the temporal resolution within which an intraspecific difference (transition) is recorded, especially given that it corresponds to the week-long period of greatest weight gain divergence. Measuring Igf-1 secretion (e.g. from the liver) and receptivity (e.g. organ-specific Igf-1 receptor levels) during this period (Hau, 2007; Cox et al., 2009; Ketterson et al., 2009) will help connect observed changes in Igf-1 to the evolution of body size.

We documented another intriguing connection between organ growth and hormonal signals. The elevated growth rate of the pancreas from 3-4 weeks in Gough mice relative to mainland mice precedes a greater separation of glucagon and glucose plasma concentrations in these strains after 4 weeks. Given that the pancreas secretes glucagon and glucagon promotes blood glucose levels, these two observations may be linked.

We also observed signatures of biochemical endophenotypes with daily precision. The divergent signatures of fuel use we documented suggest that extreme weight in Gough mice is connected to shifts in their carbohydrate and fat metabolism in the latter half of the 5^th^ postnatal week. Viewed in the context of previous studies on metabolic rate (a measurement related to RER) in mice, temporal differences in fuel source use between Gough mice and mainland mice could be related to differences in organ size (particularly that of the liver) and crosstalk between organ systems (Selman et al., 2001; Kaiyala et al., 2010; Speakman, 2013).

Overall, our results suggest that the largest wild house mice in the world evolved their unusual size principally from shifts in the accumulation and metabolism of fat during two periods of exceptional pre-pubescent growth, from remarkable growth of the liver over a week-long period, from alterations to plasma levels of hunger-inducing and growth hormones, and not from a reduction in physical activity or augmented food consumption. Similar studies of other populations hold great potential for revealing general mechanisms responsible for the island rule.

## Author contributions

MJN and BAP conceived of the study; MJN collected and analyzed the data and wrote the manuscript; BAP provided guidance on statistical analyses and edited the manuscript.

## Acknowledgements

Immunohistochemistry (IHC) was made possible by the Translational Research Initiatives in Pathology (TRIP) laboratory in the University of Wisconsin-Madison Department of Pathology and Laboratory Medicine. Anupama Singh was especially helpful in ensuring the success of the IHC. Beth Gray provided advice on the collection, fixing, and embedding of organs in preparation for IHC. Melissa Graham aided in organ dissection and histological techniques. Brian Parks and his lab members provided feedback on fat depot and liver dissection. Chris Vinyard provided guidelines for muscle dissections. We thank Michelle Parmenter for the picture of the mice in Figure 1 of the manuscript. We thank animal care staff and administrators at the UW-Madison Charmany Instructional Facility for assistance in maintaining our research mouse colony. MJN and BAP have no conflicts of interest to declare.

