## Supplementary Tables for "Phenotypic and Developmental Dissection of an Instance of the Island Rule"

**Supplementary Table S1. First instances of significant organ weight differences between Gough mice and mainland mice. T-test.**

| Organ | Female | | | Male | | |
|---|---|---|---|---|---|---|
| | Week[b] | t[a] | p-value | Week[b] | t[a] | p-value |
| **Pancreas** | 2[c] | 6.14 (24) | < 0.001 | 3 | 3.77 (22) | < 0.01 |
| **Liver** | 2[c] | 4.32 (24) | < 0.001 | 3 | 3.10 (22) | < 0.01 |
| **Mesenteric fat** | 2 | 6.91 (22) | < 0.001 | 2 | 3.20 (25) | < 0.01 |
| **Gonadal fat** | 2 | 6.72 (22) | < 0.001 | 2 | 4.96 (25) | < 0.001 |
| **Retroperitoneal fat** | 2 | 9.91 (22) | < 0.001 | 2 | 6.76 (25) | < 0.001 |
| **Subcutaneous fat** | 2 | 3.15 (22) | < 0.01 | 3[d] | 5.12 (22) | < 0.001 |
| **Brown fat** | 3[d] | 4.14 (22) | < 0.001 | 3[d] | 7.19 (22) | < 0.001 |
| **Quadriceps** | 5[e] | 5.07 (23) | < 0.001 | 5 | 4.97 (27) | < 0.001 |
| **Triceps surae** | 5[e] | 5.59 (23) | < 0.001 | 5[e] | 5.62 (27) | < 0.001 |
| **Brain** | 5[f] | 5.07 (23) | < 0.001 | 5[f] | 3.21 (27) | < 0.01 |
| **Intestine** | 2 | 13.16 (22) | < 0.001 | 2 | 11.10 (25) | < 0.001 |

[a] Degrees of freedom in parentheses.

[b] Unless labeled with a superscript, significantly different at all later time points.

[c] Significant difference at all later time points except week 3.

[d] Significant difference at all later time points except week 4.

[e] Significant difference at all later time points except week 14.

[f] Significant difference only at this time point.

| Supplementary Table S2. Planned non-orthogonal contrasts. Growth rates across weekly transitions. T-test. | | | | | | | | | | |
|---|---|---|---|---|---|---|---|---|---|---|
| Organ | Transition: week 2 to week 3[a] | | Tr: wk 3 to wk 4 | | Tr: wk 4 to wk 5 | | Tr: wk 5 to wk 6 | | Tr: wk 6 to wk 14 | |
| | t[b] | p-value* | t[b] | p-value* | t[b] | p-value* | t[b] | p-value* | t[b] | p-value* |
| **Pancreas** | 1.7 (289) | 0.096 | ***3.3 (289)*** | ***< 0.001*** | 1.3 (289) | 0.181 | 0.4 (289) | 0.680 | 0.7 (289) | 0.505 |
| **Liver** | 0.2 (292) | 0.862 | 0.9 (292) | 0.378 | ***4.5 (292)*** | ***< 0.001*** | 0.2 (292) | 0.810 | 2 (292) | 0.050 |
| **Mesenteric fat** | 1.2 (290) | 0.213 | 0.7 (290) | 0.459 | ***3.9 (290)*** | ***< 0.001*** | 1.9 (290) | 0.053 | 0.5 (290) | 0.626 |
| **Gonadal fat** | ***3.4 (291)*** | ***< 0.001*** | ***2.4 (291)*** | ***< 0.05*** | ***5.3 (291)*** | ***< 0.001*** | 1.9 (291) | 0.057 | 0.4 (291) | 0.691 |
| **Retroperitoneal fat** | ***2.5 (289)*** | ***< 0.05*** | ***3 (289)*** | ***< 0.01*** | ***5.2 (289)*** | ***< 0.001*** | 1 (289) | 0.304 | ***2.5 (289)*** | ***< 0.05*** |
| **Subcutaneous fat** | 1.8 (291) | 0.069 | 1.6 (291) | 0.115 | ***3.4 (291)*** | ***< 0.001*** | 1.8 (289) | 0.076 | 0.6 (289) | 0.522 |
| **Brown fat** | ***6.6 (292)*** | ***< 0.001*** | ***5 (292)*** | ***< 0.001*** | ***3.2 (292)*** | ***< 0.01*** | ***2 (292)*** | ***< 0.05*** | 0.4 (292) | 0.656 |
| **Quadriceps** | 1.1 (292) | 0.266 | 1.1 (292) | 0.256 | ***4 (292)*** | ***< 0.001*** | 0.4 (292) | 0.720 | 1.8 (292) | 0.076 |
| **Triceps surae** | 1.9 (292) | 0.054 | 0.7 (292) | 0.506 | ***3.7 (292)*** | ***< 0.001*** | 0.9 (292) | 0.394 | ***2.6 (292)*** | ***< 0.05*** |
| **Brain** | 1.4 (291) | 0.169 | 1.5 (291) | 0.138 | ***3.2 (291)*** | ***< 0.01*** | 1.2 (291) | 0.214 | 1.2 (291) | 0.243 |
| **Intestine** | ***2.7 (292)*** | ***< 0.01*** | ***6.1 (292)*** | ***< 0.001*** | ***5.2 (292)*** | ***< 0.001*** | 0.1 (292) | 0.886 | 0 (292) | 0.975 |

[a] Transition = Tr; week = wk.

[b] Degrees of freedom in parentheses.

* Significant results are bolded and italicized.

**Supplementary Table S3. Effects of strain, sex, and week on organ weights and length. ANOVA.**

| Organ | Strain | | Sex | | Week | | Strain by Sex Interaction | | Strain by Week Interaction | |
|---|---|---|---|---|---|---|---|---|---|---|
| | F[a] | p-value* | F[a] | p-value* | F[a] | p-value* | F[a] | p-value* | F[a] | p-value* |
| **Pancreas** | ***357.5 (1, 288)*** | ***< 0.001*** | ***11.7 (1, 288)*** | ***< 0.001*** | ***631.8 (5, 288)*** | ***< 0.001*** | 0.2 (1, 288) | 0.651 | ***13.6 (5, 288)*** | ***< 0.001*** |
| **Liver** | ***260.3 (1, 291)*** | ***< 0.001*** | ***71.1 (1, 291)*** | ***< 0.001*** | ***315 (5, 291)*** | ***< 0.001*** | 1.4 (1, 291) | 0.245 | ***12 (5, 291)*** | ***< 0.001*** |
| **Mesenteric fat** | ***895.1 (1, 289)*** | ***< 0.001*** | ***9.2 (1, 289)*** | ***< 0.01*** | ***477.5 (5, 289)*** | ***< 0.001*** | 0.0 (1, 289) | 0.861 | ***25.8 (5, 289)*** | ***< 0.001*** |
| **Gonadal fat** | ***914.4 (1, 290)*** | ***< 0.001*** | ***118.7 (1, 290)*** | ***< 0.001*** | ***263.2 (5, 290)*** | ***< 0.001*** | ***14.3 (1, 290)*** | ***< 0.001*** | ***29.6 (5, 290)*** | ***< 0.001*** |
| **Retroperitoneal fat** | ***873.6 (1, 288)*** | ***< 0.001*** | ***56.3 (1, 288)*** | ***< 0.001*** | ***122.1 (5, 288)*** | ***< 0.001*** | ***6.8 (1, 288)*** | ***< 0.01*** | ***23.2 (5, 288)*** | ***< 0.001*** |
| **Subcutaneous fat** | ***236.6 (1, 290)*** | ***< 0.001*** | ***4.8 (1, 290)*** | ***< 0.05*** | ***81.6 (5, 290)*** | ***< 0.001*** | 2.7 (1, 290) | 0.104 | ***15.6 (5, 290)*** | ***< 0.001*** |
| **Brown fat** | ***139.1 (1, 291)*** | ***< 0.001*** | ***11.8 (1, 291)*** | ***< 0.001*** | ***41.2 (5, 291)*** | ***< 0.001*** | ***5.9 (1, 291)*** | ***< 0.05*** | ***16.9 (5, 291)*** | ***< 0.001*** |
| **Quadriceps** | ***74.7 (1, 291)*** | ***< 0.001*** | ***106 (1, 291)*** | ***< 0.001*** | ***489.6 (5, 291)*** | ***< 0.001*** | 3.6 (1, 291) | 0.060 | ***13.5 (5, 291)*** | ***< 0.001*** |
| **Triceps surae** | ***37.9 (1, 291)*** | ***< 0.001*** | ***77.4 (1, 291)*** | ***< 0.001*** | ***549.1 (5, 291)*** | ***< 0.001*** | 0.0 (1, 291) | 0.968 | ***10.9 (5, 291)*** | ***< 0.001*** |
| **Brain** | ***18 (1, 290)*** | ***< 0.001*** | ***15.4 (1, 290)*** | ***< 0.001*** | ***52.7 (5, 290)*** | ***< 0.001*** | 0.3 (1, 290) | 0.345 | ***3 (5, 290)*** | ***< 0.05*** |
| **Instestine** | ***3087 (1, 291)*** | ***< 0.001*** | ***15.8 (1, 291)*** | ***< 0.001*** | ***421.1 (5, 291)*** | ***< 0.001*** | ***4.4 (1, 291)*** | ***< 0.05*** | ***81 (5, 291)*** | ***< 0.001*** |

[a] Degrees of freedom in parentheses.

* Significant results are bolded and italicized.

**Supplementary Table S4: Select sex-specific organ weights and length. Mean and Standard Deviation (SD)**

| Organ | Week 2 | | | | Week 3 | | | |
|---|---|---|---|---|---|---|---|---|
| | Female | | Male | | Female | | Male | |
| | Mainland | Gough | Mainland | Gough | Mainland | Gough | Mainland | Gough |
| | Mean (SD) | Mean (SD) | Mean (SD) | Mean (SD) | Mean (SD) | Mean (SD) | Mean (SD) | Mean (SD) |
| Gonadal fat (grams) | 0.003 (0.003) | 0.012 (0.004) | 0.002 (0.002) | 0.032 (0.021) | 0.005 (0.002) | 0.052 (0.051) | 0.004 (0.003) | 0.109 (0.051) |
| Retroperitoneal fat (grams) | 0.004 (0.003) | 0.021 (0.005) | 0.004 (0.003) | 0.023 (0.009) | 0.004 (0.002) | 0.035 (0.022) | 0.004 (0.002) | 0.039 (0.019) |
| Brown fat (grams) | 0.046 (0.008) | 0.045 (0.007) | 0.048 (0.007) | 0.041 (0.010) | 0.038 (0.004) | 0.063 (0.021) | 0.036 (0.005) | 0.006 (0.101) |
| Intestine (cm) | 18.6 (0.7) | 23.7 (1.2) | 18.8 (0.6) | 23.3 (1.3) | 22.6 (1.0) | 28.2 (2.4) | 22.3 (0.8) | 30.0 (2.4) |

| Organ | Week 4 | | | | Week 5 | | | |
|---|---|---|---|---|---|---|---|---|
| | Female | | Male | | Female | | Male | |
| | Mainland | Gough | Mainland | Gough | Mainland | Gough | Mainland | Gough |
| | Mean (SD) | Mean (SD) | Mean (SD) | Mean (SD) | Mean (SD) | Mean (SD) | Mean (SD) | Mean (SD) |
| Gonadal fat (grams) | 0.010 (0.006) | 0.033 (0.021) | 0.024 (0.010) | 0.123 (0.037) | 0.022 (0.005) | 0.139 (0.054) | 0.047 (0.017) | 0.291 (0.062) |
| Retroperitoneal fat (grams) | 0.005 (0.004) | 0.022 (0.012) | 0.010 (0.004) | 0.029 (0.009) | 0.010 (0.003) | 0.051 (0.019) | 0.015 (0.005) | 0.103 (0.040) |
| Brown fat (grams) | 0.042 (0.011) | 0.048 (0.009) | 0.050 (0.009) | 0.049 (0.006) | 0.048 (0.006) | 0.072 (0.016) | 0.061 (0.012) | 0.073 (0.013) |
| Intestine (cm) | 24.3 (1.6) | 34.8 (1.6) | 24.3 (0.9) | 35.8 (1.7) | 24.8 (1.1) | 38.8 (2.1) | 25.6 (1.1) | 40.4 (2.8) |

| Organ | Week 6 | | | | Week 14 | | | |
|---|---|---|---|---|---|---|---|---|
| | Female | | Male | | Female | | Male | |
| | Mainland | Gough | Mainland | Gough | Mainland | Gough | Mainland | Gough |
| | Mean (SD) | Mean (SD) | Mean (SD) | Mean (SD) | Mean (SD) | Mean (SD) | Mean (SD) | Mean (SD) |
| Gonadal fat (grams) | 0.037 (0.012) | 0.274 (0.101) | 0.091 (0.020) | 0.410 (0.100) | 0.070 (0.028) | 0.421 (0.187) | 0.128 (0.043) | 0.444 (0.097) |
| Retroperitoneal fat (grams) | 0.015 (0.005) | 0.083 (0.027) | 0.025 (0.005) | 0.131 (0.037) | 0.017 (0.008) | 0.114 (0.053) | 0.030 (0.016) | 0.171 (0.050) |
| Brown fat (grams) | 0.051 (0.012) | 0.082 (0.020) | 0.064 (0.006) | 0.087 (0.016) | 0.040 (0.007) | 0.069 (0.014) | 0.053 (0.012) | 0.074 (0.011) |
| Intestine (cm) | 24.4 (1.4) | 38.7 (2.4) | 25.8 (1.4) | 40.7 (2.4) | 26.5 (2.2) | 40.4 (2.2) | 26.5 (1.4) | 41.7 (1.4) |

**Supplementary Table S5. Effects of strain and week on plasma analyte concentrations. ANOVA.**

| Plasma analyte | Female | | | | | | Male | | | | | |
|---|---|---|---|---|---|---|---|---|---|---|---|---|
| | Strain | | Week | | Strain by Week Interaction | | Strain | | Week | | Strain by Week Interaction | |
| | F[a] | p-value* | F[a] | p-value* | F[a] | p-value* | F[a] | p-value* | F[a] | p-value* | F[a] | p-value* |
| **Ghrelin** | ***97.7 (1, 94)*** | ***< 0.001*** | ***11 (5, 94)*** | ***< 0.001*** | ***7.0 (5, 94)*** | ***< 0.001*** | ***201.7 (1, 93)*** | ***< 0.001*** | ***21.8 (5, 93)*** | ***< 0.001*** | ***8.4 (5, 93)*** | ***< 0.001*** |
| **Leptin** | ***244.1 (1, 94)*** | ***< 0.001*** | ***12.3 (5, 94)*** | ***< 0.001*** | ***25.2 (5, 94)*** | ***< 0.001*** | ***118.6 (1, 91)*** | ***< 0.001*** | ***9.7 (5, 91)*** | ***< 0.001*** | ***6.1 (5, 91)*** | ***< 0.001*** |
| **Glucagon** | ***99.5 (1, 95)*** | ***< 0.001*** | ***6.6 (5, 95)*** | ***< 0.001*** | ***4.8 (5, 95)*** | ***< 0.01*** | ***20.2 (1, 91)*** | ***< 0.001*** | ***3.7 (5, 91)*** | ***< 0.01*** | 1 (5, 91) | 0.43 |
| **Glucose** | ***63.1 (1, 135)*** | ***< 0.001*** | ***24 (5, 135)*** | ***< 0.001*** | 1.3 (5, 135) | 0.258 | ***129.7 (1, 121)*** | ***< 0.001*** | ***52.3 (5, 121)*** | ***< 0.001*** | ***7.4 (5, 121)*** | ***< 0.001*** |
| **Adiponectin** | ***27.9 (1, 90)*** | ***< 0.001*** | ***51.9 (5, 90)*** | ***< 0.001*** | ***3.7 (5, 90)*** | ***< 0.01*** | 4.1 (1, 90) | 0.05 | ***44 (5, 90)*** | ***< 0.001*** | ***5.2 (5, 90)*** | ***< 0.001*** |
| **IGF-1** | ***31.3 (1, 98)*** | ***< 0.001*** | ***4.7 (5, 98)*** | ***< 0.01*** | ***6.4 (5, 98)*** | ***< 0.001*** | 0.3 (1, 98) | 0.582 | ***7.6 (5, 98)*** | ***< 0.001*** | ***7.2 (5, 98)*** | ***< 0.001*** |
| **Insulin** | 0.1 (1, 133) | 0.713 | ***17.7 (5, 133)*** | ***< 0.001*** | 1.4 (5, 133) | 0.223 | 0.7 (1, 121) | 0.418 | ***10.2 (5, 121)*** | ***< 0.001*** | 0.5 (5, 121) | 0.776 |
| **Free fatty acid** | ***43 (1, 103)*** | ***< 0.001*** | ***6.5 (5, 103)*** | ***< 0.001*** | 1.8 (5, 103) | 0.13 | 0.01 (1, 91) | 0.921 | ***7.3 (5, 91)*** | ***< 0.001*** | 0.7 (5, 91) | 0.629 |

[a] Degrees of freedom in parentheses.

* Significant results are bolded and italicized.

**Supplementary Table S6. Temporal differences in Respiratory Exchange Ratio between Gough mice and mainland mice. T-test.**

| Rest/Activity Classification | Female | | | | | | | | Male | | | | | | | |
|---|---|---|---|---|---|---|---|---|---|---|---|---|---|---|---|---|
| | Day 29 | | | | Day 34 | | | | Day 29 | | | | Day 34 | | | |
| | Mean (Standard deviation) | | $t^a$ | p-value* | Mean (Standard deviation) | | $t^a$ | p-value* | Mean (Standard deviation) | | $t^a$ | p-value* | Mean (Standard deviation) | | $t^a$ | p-value* |
| | Mainland | Gough | | | Mainland | Gough | | | Mainland | Gough | | | Mainland | Gough | | |
| **Circadian Rest** | 0.85 (0.06) | 0.80 (0.03) | 2.1 (13) | 0.05 | ***0.79 (0.02)*** | ***0.88 (0.02)*** | ***7.9 (15)*** | ***< 0.001*** | 0.77 (0.05) | 0.79 (0.03) | 0.9 (12) | 0.39 | ***0.78 (0.03)*** | ***0.89 (0.04)*** | ***6.5 (14)*** | ***< 0.001*** |
| **Rest - No Activity** | ***0.90 (0.04)*** | ***0.84 (0.04)*** | ***3.1 (13)*** | ***< 0.01*** | ***0.83 (0.02)*** | ***0.92 (0.03)*** | ***6.5 (15)*** | ***< 0.001*** | 0.84 (0.06) | 0.82 (0.05) | 0.7 (12) | 0.47 | 0.86 (0.06) | 0.89 (0.04) | 1.4 (14) | 0.18 |
| **NonWheel Activity** | 0.94 (0.06) | 0.88 (0.04) | 2.2 (13) | 0.05 | 0.91 (0.03) | 0.93 (0.03) | 1.2 (15) | 0.25 | 0.89 (0.05) | 0.85 (0.03) | 1.6 (12) | 0.14 | 0.93 (0.03) | 0.95 (0.02) | 1.8 (14) | 0.09 |
| **Wheel - Long** | ***1.02 (0.01)*** | ***0.95 (0.04)*** | ***3.5 (12)*** | ***< 0.01*** | 0.96 (0.04) | 0.99 (0.03) | 1.7 (15) | 0.11 | 0.97 (0.07) | 0.92 (0.03) | 1.5 (12) | 0.15 | 0.98 (0.03) | 1 (0.01) | 1.3 (14) | 0.23 |
| **Wheel - Spurt** | ***1.02 (0.02)*** | ***0.92 (0.02)*** | ***4.5 (13)*** | ***< 0.001*** | 1.01 (0.03) | 0.99 (0.02) | 1.1 (15) | 0.31 | 0.98 (0.08) | 0.93 (0.07) | 1.6 (12) | 0.14 | 1.01 (0.04) | 1 (0.05) | 0.5 (14) | 0.63 |

[a] Degrees of freedom in parentheses.

* Significant results are bolded and italicized. Mean RER higher in mainland mice labeled in green. Mean RER higher in Gough mice labeled in blue.

| Supplementary Table S7. Effects of strain and day on Respiratory Exchange Ratio. ANOVA. | | | | | | | | | | | | |
|---|---|---|---|---|---|---|---|---|---|---|---|---|
| Rest/Activity Classification | Female | | | | | | Male | | | | | |
| | Strain | | Day | | Strain by Day Interaction | | Strain | | Day | | Strain by Day Interaction | |
| | F[a] | p-value* | F[a] | p-value* | F[a] | p-value* | F[a] | p-value* | F[a] | p-value* | F[a] | p-value* |
| **Circadian Rest** | ***24,9 (1, 94)*** | ***< 0.001*** | ***5.0 (1, 94)*** | ***< 0.05*** | ***28.8 (1, 94)*** | ***< 0.001*** | ***47.1 (1, 90)*** | ***< 0.001*** | 1.0 (1, 90) | 0.314 | ***19.6 (1, 90)*** | ***< 0.001*** |
| **Rest - No Activity** | ***31.2 (1, 92)*** | ***< 0.001*** | ***6.2 (1, 92)*** | ***< 0.05*** | ***33.5 (1, 92)*** | ***< 0.001*** | 2.8 (1, 85) | 0.101 | 0.0 (1, 85) | 0.980 | ***3.0 (1, 85)*** | ***< 0.1*** |
| **NonWheel Activity** | ***8.4 (1, 94)*** | ***< 0.01*** | 0.8 (1, 94) | 0.378 | ***8.3 (1, 94)*** | ***< 0.01*** | ***16.4 (1, 90)*** | ***< 0.001*** | 0.0 (1, 90) | 0.871 | ***15.1 (1, 90)*** | ***< 0.001*** |
| **Wheel - Long** | ***15.6 (1, 92)*** | ***< 0.001*** | ***6.0 (1, 92)*** | ***< 0.05*** | ***15.2 (1, 92)*** | ***< 0.001*** | ***18.4 (1, 90)*** | ***< 0.001*** | 0.6 (1, 90) | 0.429 | ***16.5 (1, 90)*** | ***< 0.001*** |
| **Wheel - Spurt** | 1.9 (1, 94) | 0.167 | 0.1 (1, 94) | 0.751 | 1.5 (1, 94) | 0.223 | ***8.0 (1, 90)*** | ***< 0.001*** | 0.7 (1, 90) | 0.396 | ***6.7 (1, 90)*** | ***< 0.05*** |

[a] Degrees of freedom in parentheses.

* Significant results are bolded and italicized.

**Supplementary Table S8. Effects of strain on food and water consumption. T-test.**

| Day | Food consumption (grams) | | | | | | | | Water consumption (ml) | | | | | | | |
|---|---|---|---|---|---|---|---|---|---|---|---|---|---|---|---|---|
| | Female | | | | Male | | | | Female | | | | Male | | | |
| | mean (standard deviation) | | t[a] | p-value* | mean (standard deviation) | | t[a] | p-value* | mean (standard deviation) | | t[a] | p-value* | mean (standard deviation) | | t[a] | p-value* |
| | Mainland | Gough | | | Mainland | Gough | | | Mainland | Gough | | | Mainland | Gough | | |
| **29** | 2.8 (0.3) | 2.4 (0.4) | 2.5 (13) | 0.029 | ***3.1 (0.3)*** | ***2.5 (0.4)*** | ***3.5 (12)*** | ***< 0.01*** | ***2.3 (0.2)*** | ***4.1 (0.8)*** | ***5.9 (13)*** | ***< 0.001*** | ***2.6 (0.2)*** | ***3.9 (0.9)*** | ***3.5 (11)*** | ***< 0.01*** |
| **30** | 3.0 (0.3) | 2.6 (0.3) | 2.2 (13) | 0.043 | ***3.6 (0.2)*** | ***2.8 (0.4)*** | ***4.6 (14)*** | ***< 0.001*** | ***2.4 (0.2)*** | ***4.1 (0.9)*** | ***4.8 (13)*** | ***< 0.001*** | ***2.6 (0.2)*** | ***4.1 (0.8)*** | ***5.0 (13)*** | ***< 0.001*** |
| **31** | 3.0 (0.2) | 3.0 (0.4) | 0.06 (15) | 0.952 | ***3.7 (0.3)*** | ***3.2 (0.2)*** | ***4.3 (14)*** | ***< 0.001*** | ***2.6 (0.3)*** | ***4.5 (0.9)*** | ***6.3 (15)*** | ***< 0.001*** | ***2.7 (0.2)*** | ***4.4 (1.0)*** | ***4.8 (13)*** | ***< 0.001*** |
| **32** | 3.2 (0.3) | 3.2 (0.6) | 0.21 (15) | 0.840 | 3.8 (0.2) | 3.5 (0.5) | 1.2 (14) | 0.237 | ***2.8 (0.2)*** | ***4.6 (1.1)*** | ***5.2 (15)*** | ***< 0.001*** | ***2.8 (0.1)*** | ***4.8 (1.0)*** | ***5.7 (13)*** | ***< 0.001*** |
| **33** | 3.5 (0.2) | 3.5 (0.5) | 0.37 (15) | 0.717 | 3.8 (0.3) | 4.1 (0.5) | 1.2 (14) | 0.234 | ***2.8 (0.5)*** | ***4.8 (1.1)*** | ***5.1 (15)*** | ***< 0.001*** | ***2.7 (0.2)*** | ***5.0 (1.2)*** | ***5.6 (13)*** | ***< 0.001*** |
| **34** | 3.4 (0.5) | 3.7 (0.5) | 1.2 (15) | 0.241 | 3.9 (0.3) | 4.4 (0.8) | 1.6 (14) | 0.138 | ***2.6 (0.4)*** | ***4.9 (1.1)*** | ***6.1 (15)*** | ***< 0.001*** | ***2.8 (0.2)*** | ***5.2 (1.1)*** | ***5.9 (13)*** | ***< 0.001*** |

[a] Degrees of freedom in parentheses.

* Significant results are bolded and italicized.
