## Supplementary Figures for "Phenotypic and Developmental Dissection of an Instance of the Island Rule"

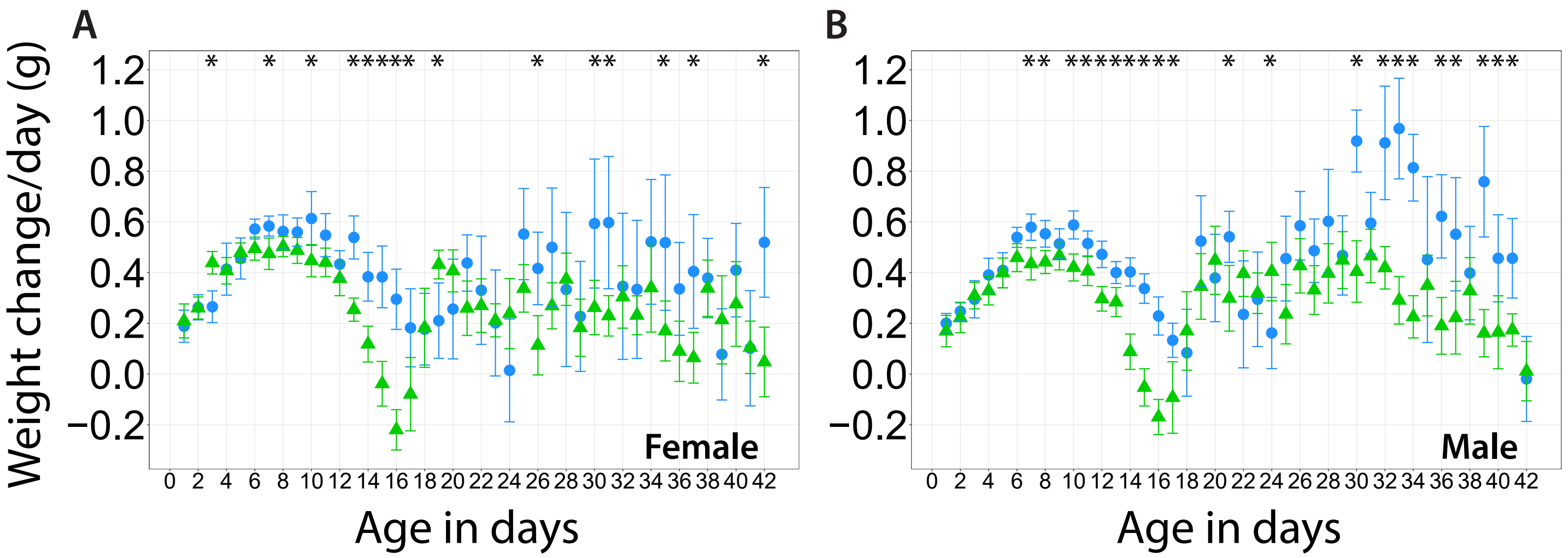


**Supplementary Figure 1.** Day-to-day weight change. Color and shape designations are consistent across panels: Blue circles, Gough; green triangles, Mainland. Asterisks mark significant differences; error bars are ± 2 x standard error of the means.

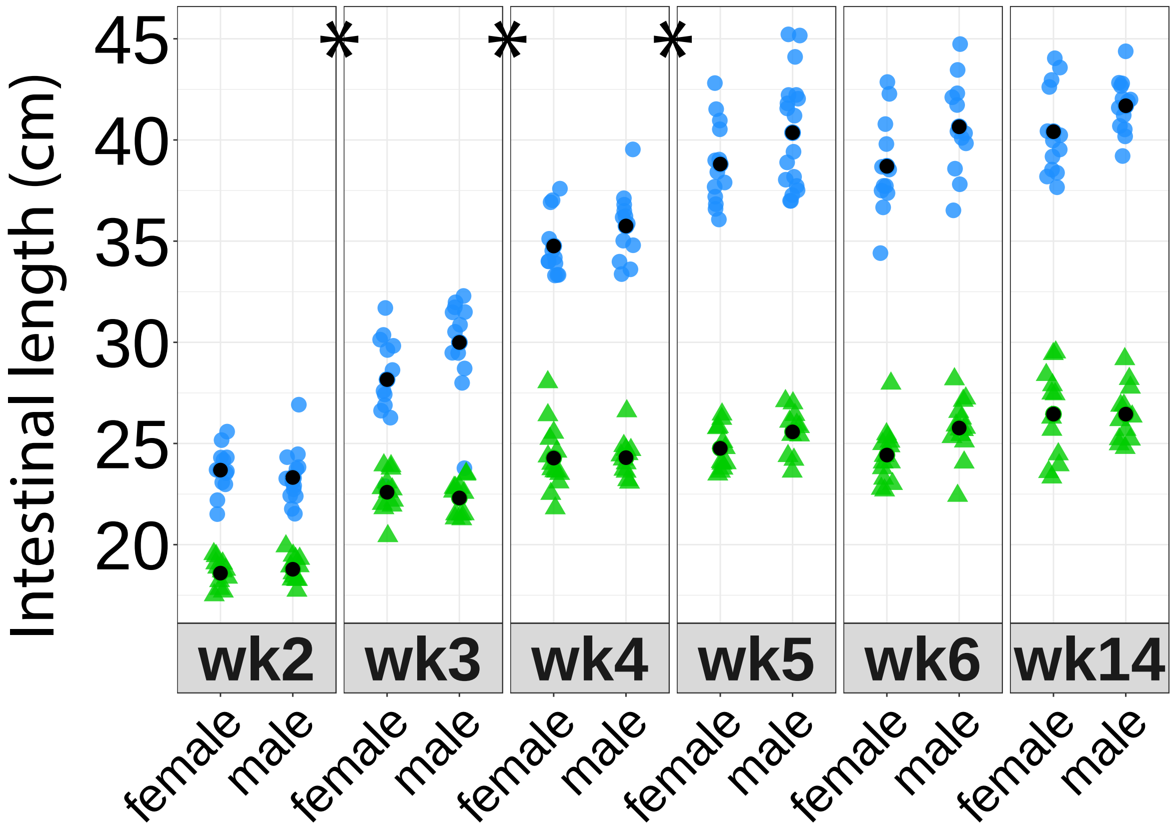


**Supplementary Figure 2.** Weekly intestinal length. Asterisks mark weekly transitions when the rate of intestinal lengthening is greater in Gough mice than Mainland mice. Blue circles signify measurements from Gough mice; green triangles signify measurements from Mainland mice; black circles signify mean intestinal lengths.

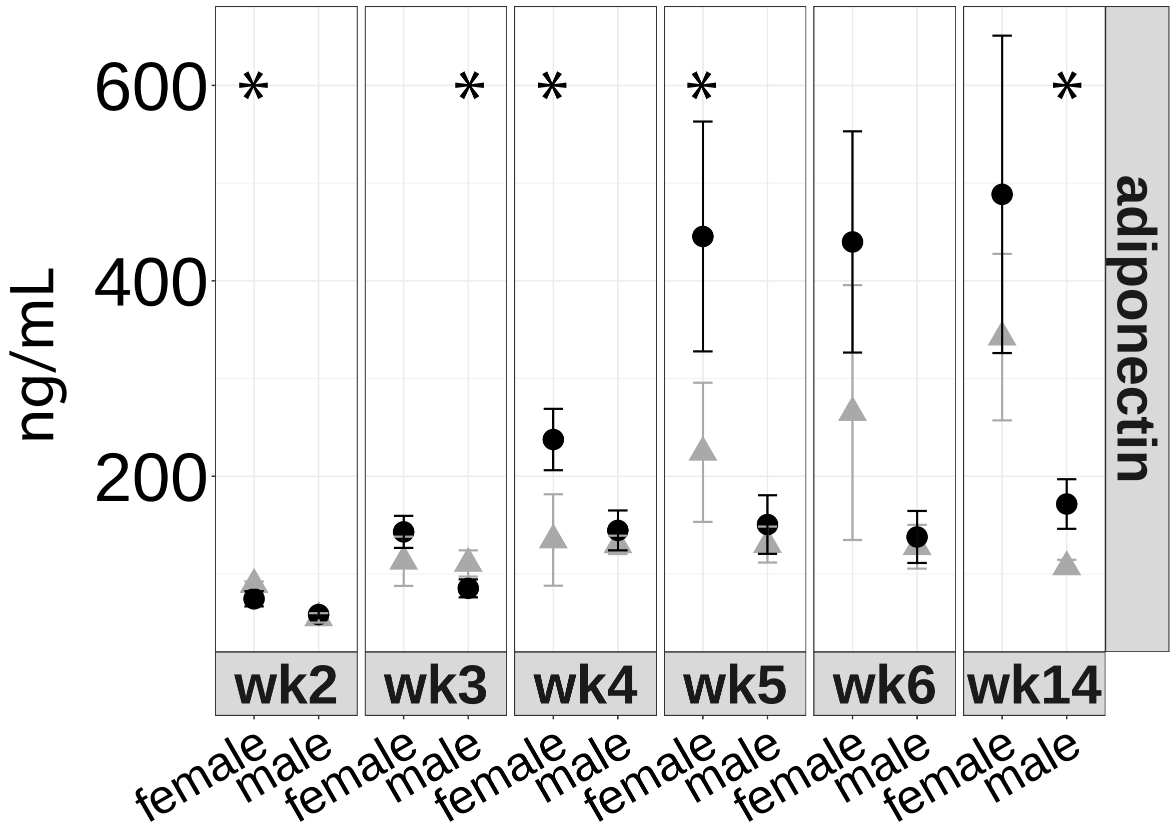


**Supplementary Figure 3.** Weekly fasting mean adipocyte concentration. Black circles signify measurements from Gough mice; gray triangles signify measurements from Mainland mice. Error bars are ± 2 x standard error of the means and asterisks mark significant differences.

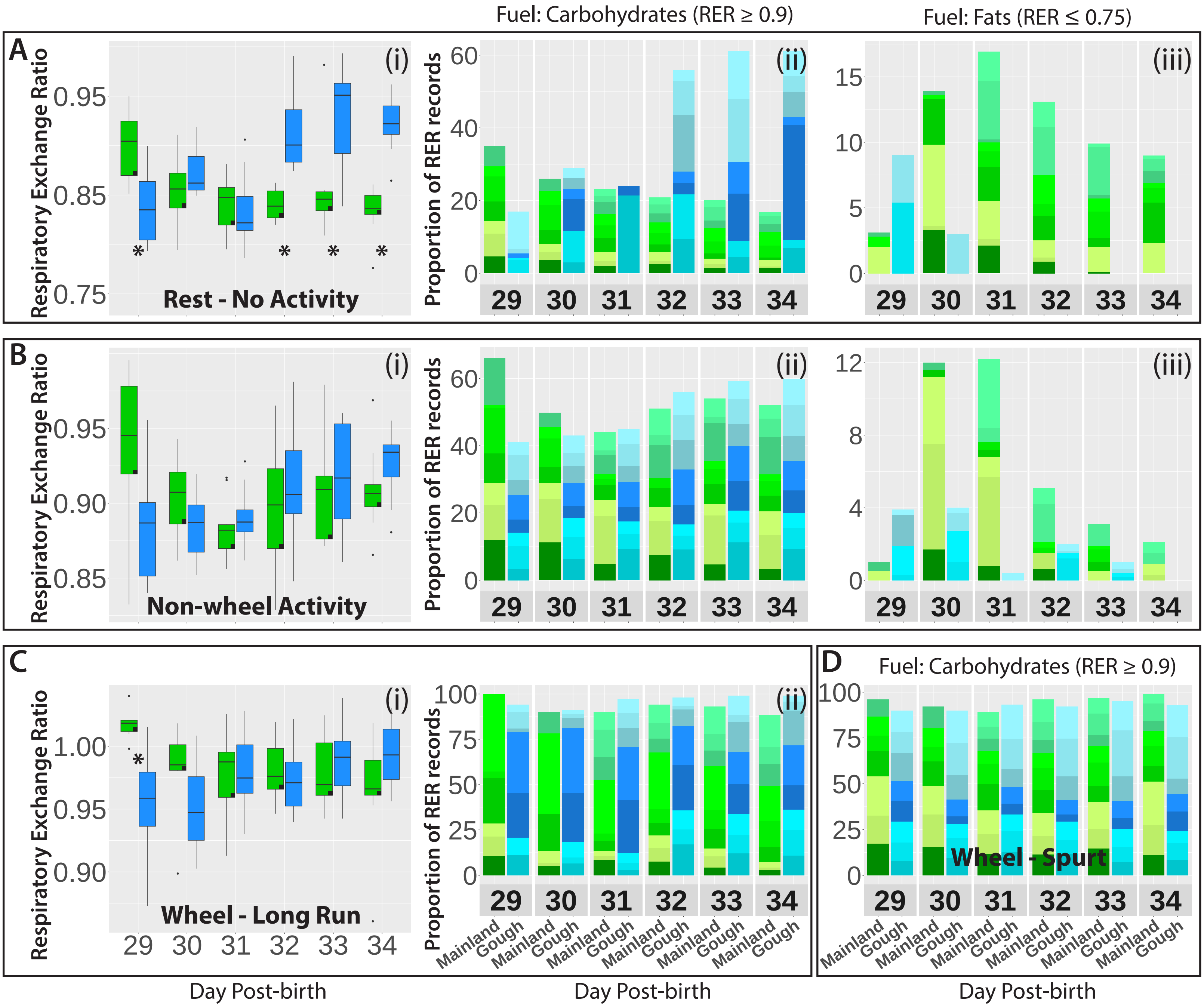


**Supplementary Figure 4.** Comparison of respiratory exchange ratios (RER) in Females. (A, B, C, D) Different rest-activity classes from which RERs were taken. Box plots in panels labeled (i) treat the mean RER within a rest-activity class for a mouse as a data point and pooled strain-specific mouse means across days. Blue signifies measurements from Gough mice; green signifies measurements from Mainland mice. When present, black boxes in lower right of box plots also indicate Mainland strain. Asterisks mark significant differences. All panels labeled (ii) and panel (D) show the proportion of RERs for a given rest-activity class that are categorically indicative of energy production via carbohydrates (RER record ≥ 0.9). All panels labeled (iii) show the proportion of RERs for a given rest-activity class that are categorically indicative of energy production via fats (RER record ≤ 0.75). The color hues in panels labeled (ii), (iii), and (D) signify, generally, the strain (green hues = Mainland; blue hues = Gough) and specifically the mouse (each hue representing contributions from a single mouse) from which an RER record was obtained. It is important to consider how the ranges of RER change both across time (days 29-24) and from restful to strenuous behavior (moving from top to bottom in the Figure). As the rest-activity classes move from restful (Rest - No Activity) to strenuous (Wheel-Long Run), the range of average RER becomes more constrained from day 29 to day 34. In the Wheel-Long Run class (Ci), most of the average RER ranges are positioned above 0.9. The generally high average RER values associated with increasingly strenuous activity - Wheel-Long Run (Ci) and Wheel-Spurt (D) likely do not represent an accurate picture of metabolic fuel source in the strains; rather, they indicate a shift towards the exhalation of more $CO_2$ to offset anaerobic acidification of the blood (Seidenberg and Beutler 2008). This observation validates our approach to divide average pooled RER values into different rest-activity classes and indicates that strain differences in RER patterns in more restful categories offer insight into divergent metabolic processes that accompany extreme weight gain in Gough mice.

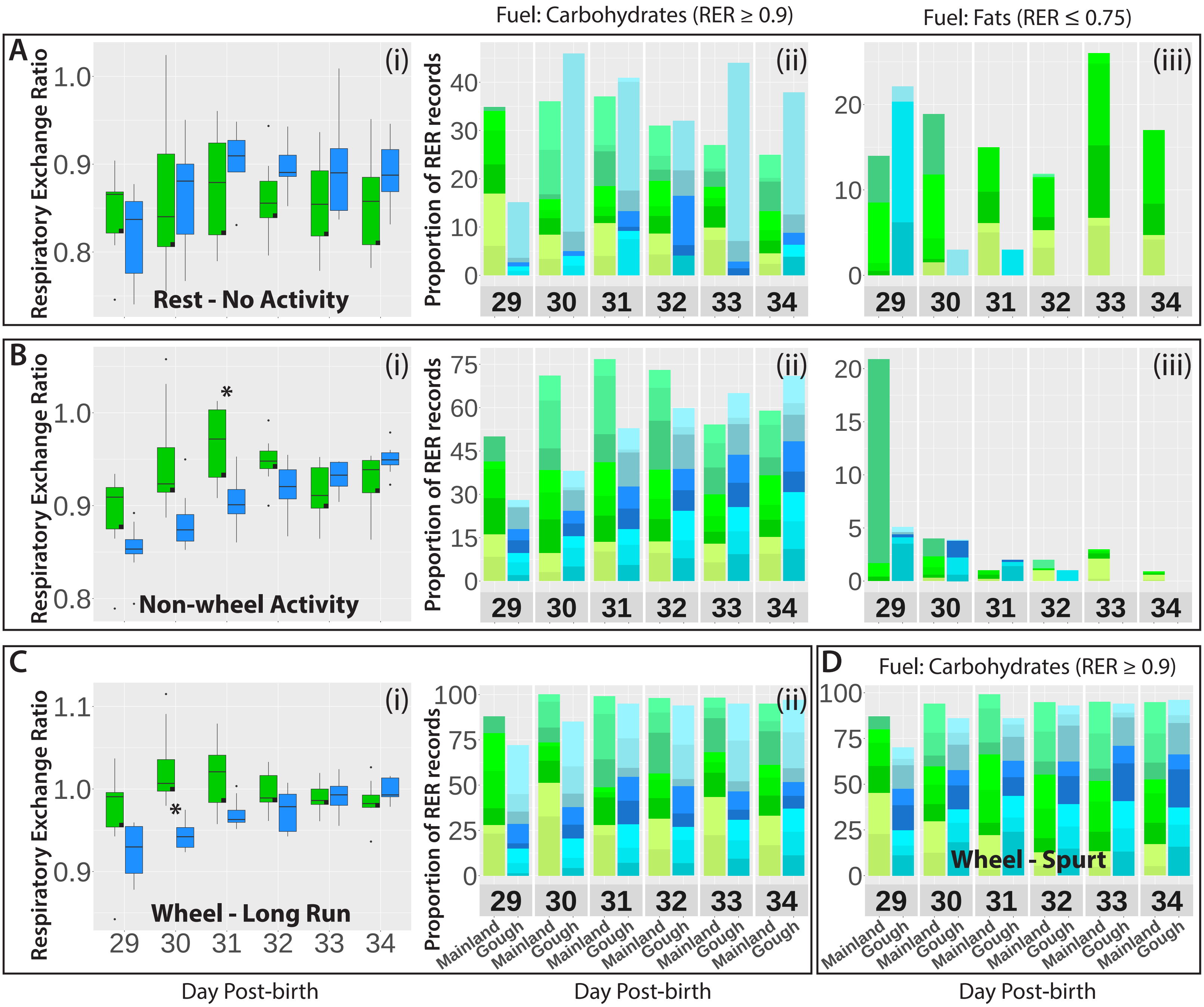


**Supplementary Figure 5.** Comparison of respiratory exchange ratios (RER) in Males. (A, B, C, D) Different rest-activity classes from which RERs were taken. Box plots in panels labeled (i) treat the mean RER within a rest-activity class for a mouse as a data point and pooled strain-specific mouse means across days. Blue signifies measurements from Gough mice; green signifies measurements from Mainland mice. When present, black boxes in lower right of box plots also indicate Mainland strain. Asterisks mark significant differences. All panels labeled (ii) and panel (D) show the proportion of RERs for a given rest-activity class that are categorically indicative of energy production via carbohydrates (RER record ≥ 0.9). All panels labeled (iii) show the proportion of RERs for a given rest-activity class that are categorically indicative of energy production via fats (RER record ≤ 0.75). The color hues in panels labeled (ii), (iii), and (D) signify, generally, the strain (green hues = Mainland; blue hues = Gough) and specifically the mouse (each hue representing contributions from a single mouse) from which an RER record was obtained. It is important to consider how the ranges of RER change both across time (days 29-24) and from restful to strenuous behavior (moving from top to bottom in the Figure). As the rest-activity classes move from restful (Rest - No Activity) to strenuous (Wheel-Long Run), the range of average RER becomes more constrained from day 29 to day 34. In the Wheel-Long Run class (Ci), most of the average RER ranges are positioned above 0.9. The generally high average RER values associated with increasingly strenuous activity - Wheel-Long Run (Ci) and Wheel-Spurt (D) likely do not represent an accurate picture of metabolic fuel source in the strains; rather, they indicate a shift towards the exhalation of more $CO_2$ to offset anaerobic acidification of the blood (Seidenberg and Beutler 2008). This observation validates our approach to divide average pooled RER values into different rest-activity classes and indicates that strain differences in RER patterns in more restful categories offer insight into divergent metabolic processes that accompany extreme weight gain in Gough mice.
